# Heterochronic parabiosis reprograms the mouse brain transcriptome by shifting aging signatures in multiple cell types

**DOI:** 10.1101/2022.01.27.477911

**Authors:** Methodios Ximerakis, Kristina M. Holton, Richard M. Giadone, Ceren Ozek, Monika Saxena, Samara Santiago, Xian Adiconis, Danielle Dionne, Lan Nguyen, Kavya M. Shah, Jill M. Goldstein, Caterina Gasperini, Scott L. Lipnick, Sean K. Simmons, Sean M. Buchanan, Amy J. Wagers, Aviv Regev, Joshua Z. Levin, Lee L. Rubin

## Abstract

Aging is a complex process involving transcriptomic changes associated with deterioration across multiple tissues and organs, including the brain. Recent studies using heterochronic parabiosis have shown that various aspects of aging-associated decline are modifiable or even reversible. To better understand how this occurs, we performed single-cell transcriptomic profiling of young and old mouse brains following parabiosis. For each cell type, we catalogued alterations in gene expression, molecular pathways, transcriptional networks, ligand-receptor interactions, and senescence status. Our analyses identified gene signatures demonstrating that heterochronic parabiosis regulates several hallmarks of aging in a cell-type-specific manner. Brain endothelial cells were found to be especially malleable to this intervention, exhibiting dynamic transcriptional changes that affect vascular structure and function. These findings suggest novel strategies for slowing deterioration and driving regeneration in the aging brain through approaches that do not rely on disease-specific mechanisms or actions of individual circulating factors.

## Introduction

Aging is a complicated multifactorial process that is far from being completely understood. However, the recent characterization of highly conserved and interconnected hallmarks of aging has unified and advanced aging research^1^. This progress has ultimately contributed to a better understanding of how tissues and organs change with age. In particular, the brain has been shown to undergo a variety of changes at the molecular, cellular, and system levels throughout organismal aging. These changes include defects in cellular respiration and protein synthesis, increased oxidative stress, decreased neurotransmission through loss of synapses and neuronal atrophy, reduced myelination, reduction in the regenerative capacity of neural stem cells, increased inflammation, and the impairment of blood vessels and blood flow^2,3^, many of which also accompany several late-onset neurodegenerative diseases^4,5^. This is significant given that a series of recent observations demonstrate that several aspects of aging can be delayed or even reversed by a variety of interventions, including exercise^6,7^, caloric restriction^8^, elimination of senescent cells^9,10^, administration of rapamycin^11^ or metformin^12^, transient cell reprogramming^13^, and young bone marrow transplantation^14^.

One of the most robust methods of improving the function of aging tissues is that of heterochronic parabiosis, a surgical procedure whereby young and old mice are joined together so that they share a common circulatory system^15,16^. A number of seminal publications have led to the surprising conclusion that exposure of old mice to the young circulatory environment can improve the function of various tissues and organs^17–23^, including the central nervous system (CNS)^24–28^. In the CNS specifically, studies from our lab^27,29^ and others^26,30–32^ have shown that circulating factors in young blood can stimulate functional improvement in the aged or diseased brain, including improvements in neurogenesis, synapse number, neural activity and behavior^24,26–28^. Our work has also demonstrated that blood-borne factors can stimulate a global regrowth of healthy brain vasculature with improved blood flow in old mice^27,29^. Conversely, it has been shown that systemic factors in old blood drive aging phenotypes in young tissues^33,34^, including the brain, whereby inhibition of neurogenesis and impairment of cognitive function have been described^24,34,35^. It has also been reported that the dilution of old blood plasma alone is sufficient to trigger a multi-tissue rejuvenation process in old mice^36,37^, while the dilution of the young blood plasma alone does not induce aging acceleration processes in young mice^36^.

While parabiosis has a profound effect on brain function, its molecular underpinnings remain to be fully elucidated. Recent efforts have employed proteomics to measure age-related changes in serum blood proteins^38^, primarily based on the hypothesis that function-improving factors decline with age. While this is a plausible and useful approach^27,30,31,39–41^, it ignores non-protein factors (blood cells, exosomes, lipids) ^25^ and factors that either do not change or change in unexpected directions with aging.

Following our previously-published work employing single cell RNA sequencing (scRNA-seq) to comprehensively describe changes that occur in the brain during aging^42^, we quantified changes in the transcriptome of young and old mouse brains subjected to parabiosis. The comprehensive single-cell datasets generated in this work allowed us to detect brain cell types whose transcriptional states are affected by parabiosis, as determined by their differentially expressed genes. In this characterization, we also identified inter/intracellular molecular pathways, gene regulatory networks, cell-cell interactions, and senescence state changes that are associated with rejuvenation and/or aging acceleration. Endothelial cells were found to be highly affected by parabiosis, exhibiting strong dynamic changes in their transcriptome, with several of these validated by orthogonal assays. Overall, this work shows that heterochronic parabiosis regulates manifestations of several canonical hallmarks of aging^1^, such as the altered intercellular communication and cellular senescence, by shifting aging-induced changes of the transcriptome in a cell-type-specific manner.

## Results

### Single-cell transcriptomic profiling of parabiotic brains to study rejuvenation and aging acceleration

To gain new insights into the effects of heterochronic parabiosis on the mouse brain and to assess its molecular effects on individual cell types, we employed high-throughput single-cell RNA sequencing (scRNA-seq) to examine the transcriptional profiles of young and old mouse brains following parabiosis (Fig 1a-b). We generated heterochronic pairs in which 3-4-month-old mice were joined with 20-22-month-old mice (analogous to human ages 18-20 and 65-70 years respectively)^43,44^. We also generated age-matched isochronic pairs of young and old mice as controls. All parabiotic pairs were maintained for 4-5 weeks before tissue collection and analysis. We confirmed successful parabiosis and establishment of blood cross-circulation by demonstrating the presence in both partners of heterochronic pairs of blood cells bearing congenic markers from both the aged (CD45.2^+^) and young (CD45.1^+^) parabiont, as previously described^45^(Supplementary Fig 1). We dissociated the brain tissues from parabiotic mice using our recently developed protocol^42^ and analyzed the transcriptome of 158,767 single cells, where 108,555 cells were derived from the brains of 40 parabionts [8 young isochronic (hereafter referred to as YY); 9 young heterochronic (YO); 12 old isochronic (OO); 11 old heterochronic (OY)], and 50,212 cells derived from the brains of 16 unpaired mice [8 young (YX) and 8 old (OX)] (Supplementary Fig 2-3). For the unpaired mice, we integrated and re-analyzed our recently published single cell dataset^42^ that was generated simultaneously with those of the parabionts to assist in cell type identification and gene expression analysis.

**Fig 1.**
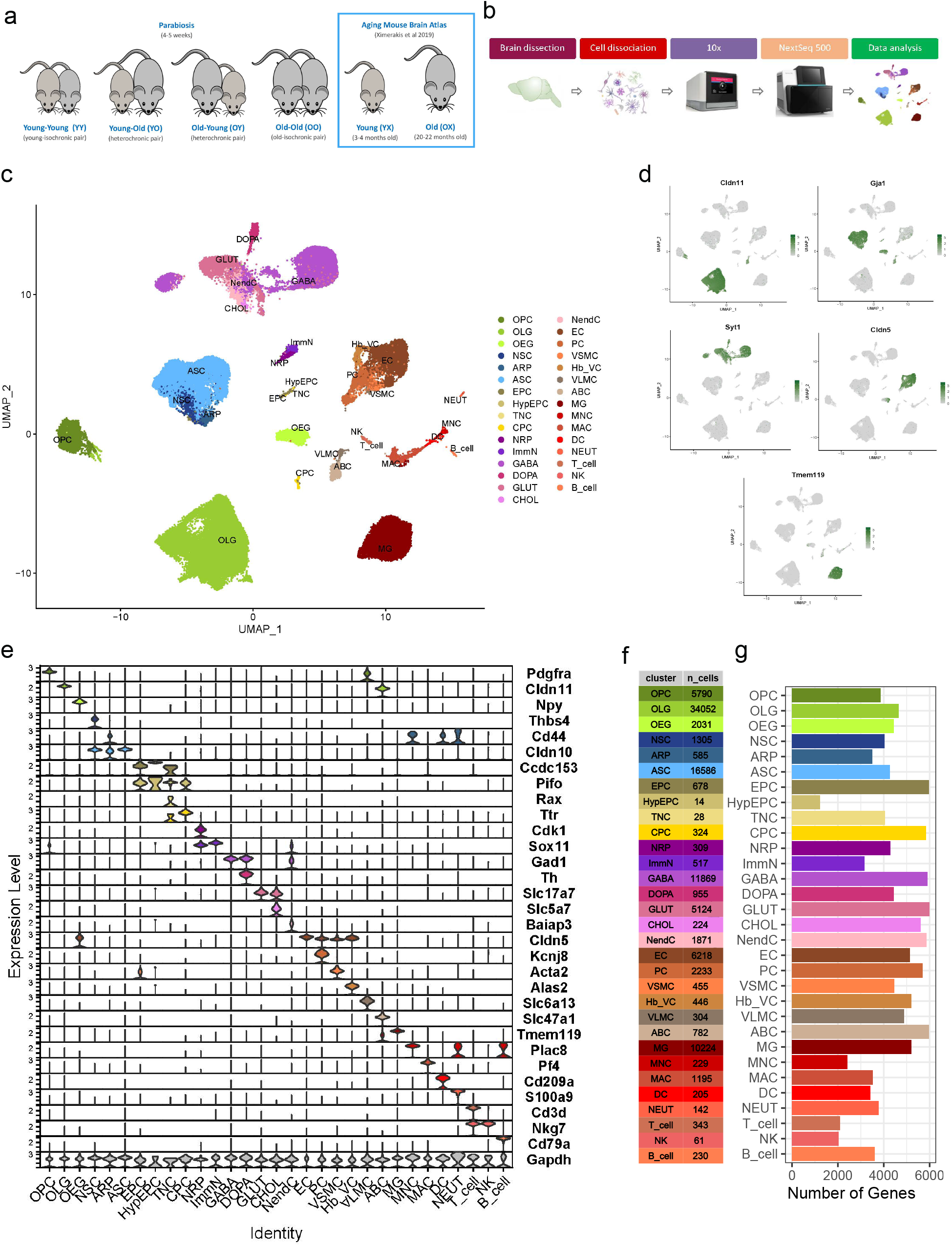
Overview of the single-cell sequencing analysis. **a**. Schematic representation of the animal types used in this study. Sequencing data from isochronic (YY and OO) and heterochronic (YO and OY) parabiosis pairs were generated and integrated with sequencing data from young (YX) and old (OX) unpaired mice from our previous work^1^. **b.** Schematic representation of the experimental workflow (see Methods for details). **c**. UMAP projection of 105,329 cells across 31 clusters derived from 34 parabionts (11 old heterochronic, 9 young heterochronic, 7 old isochronic, 7 young isochronic) and 16 unpaired mice (8 old, and 8 young). **d.** UMAP projection of 5 major cell populations showing the expression of representative well-known cell-type-specific marker genes (*Cldn11, Gja1, Syt1, Cldn5, Tmem119*). Numbers reflect the number of nCount RNA (UMI) detected for the specified gene for each cell. **e**. Violin plot showing the distribution of expression levels of well-known representative cell-type-enriched marker genes across all 31 distinct cell types. **f.** Table showing the total number of detected cells per cell type. **g.** Bar plot showing the total number of detected genes per cell type.

The scRNA-seq dataset was filtered to retain high quality reads and clearly defined clusters. Cells from all animal groups were distributed over 5 sequencing batches, with no batch effects observed that would warrant further correction (Supplementary Fig 4). We employed K-nearest neighbors clustering with Louvain community detection and removed low quality clusters resulting from debris, doublets/multiplets, and dead cells. We also excluded animals with low quality reads and employed other quality control measures as described in the Methods (Supplementary Fig 5). In total, we retained and re-clustered 105,329 cells, derived from 34 parabionts (7 YY; 9 YO; 7 OO; 11 OY), and 16 unpaired (8 YX and 8 OX) mice (Supplementary Fig 2).

### Construction of single-cell atlases reveals that cell type identity and composition are largely preserved in parabiotic brains

We first identified the cell types present by combining cells in unpaired and parabiotic brains. Using cell markers established from our previous work^42^ on young and old mouse brains, we identified 31 major cell types (Fig 1c) with distinct expression profiles: oligodendrocyte precursor cells (OPC), oligodendrocytes (OLG), olfactory ensheathing glia (OEG); neural stem cells (NSC), astrocyte-restricted precursors (ARP), astrocytes (ASC), ependymocytes (EPC), hypendymal cells (HypEPC), tanycytes (TNC), choroid plexus epithelial cells (CPC); neuronal-restricted precursors (NRP), immature neurons (ImmN), GABAergic neurons (GABA), dopaminergic neurons (DOPA), glutamatergic neurons (GLUT), cholinergic neurons (CHOL), neuroendocrine cells (NendC)^42^, endothelial cells (EC), pericytes (PC), vascular smooth muscle cells (VSMC), hemoglobin-expressing vascular cells (Hb-VC)^42^, vascular and leptomeningeal cells (VLMC), arachnoid barrier cells (ABC), microglia (MG), monocytes (MNC), macrophages (MAC), dendritic cells (DC), neutrophils (NEUT), T-cells (T_cell), natural killer cells (NK), and B-cells (B_cell) (Fig 1c, Supplementary Table 1). We then examined the major markers for each cell type to ensure robustness of their transcriptional signatures. Each cell type expressed markers that are distinct to its cellular identity, as previously characterized^42^ (Fig 1d-e).

Our unsupervised clustering also showed that the identified cell populations were represented in all batches and animal types (Supplementary Fig 4, Supplementary Fig 6a), indicating that cell identity is preserved in parabiotic mice. We found that the proportions of each cell type represented in each group were only slightly smaller for the isochronic brains, which may be due to the smaller number of contributing animals (Supplementary Fig 7). We quantified the difference in the number of cells per each cell population by comparing the animal types using ANOVA with a threshold of p ≤ 0.05, confirming that parabiosis does not significantly alter the number of cells per cell population between animal types with the exception of DOPA in young heterochronic versus young unpaired mice (p=0.002) (Fig 1f, Supplementary Fig 7, Supplementary Table 2). However, data analysis of cell proportion changes should be cautiously considered in single cell sequencing studies, especially when tissue dissociation methods are used. Across the different types of cells, we observed the largest number of detected genes in EPC, CPC, GABA, GLUT, NendC, and ABC (Fig 1g).

For further investigation, we performed high resolution subclustering analysis to uncover the heterogeneity of all identified cell types. To reduce the effects of drastically different cell identities, we first grouped the identified cells into five distinct classes (Oligodendrocyte lineage and OEG, Astro-ependymal cells and NSC, Neuronal lineage, Vasculature cells, and Immune cells) based on their expression profile, lineage, function, and anatomical organization. We delineated 75 distinct cell populations in accordance with the literature and our previous results^42^ (Fig 2a-e). However, with a greater number of animals and groups than our previous study^42^, more heterogeneity was revealed, leading to higher granularity in certain cell types, such as GABA, GLUT, NendC, EC, and MG. As with the major cell populations, all the identified subpopulations were represented in each animal type (Supplementary Fig 6b).

**Fig 2.**
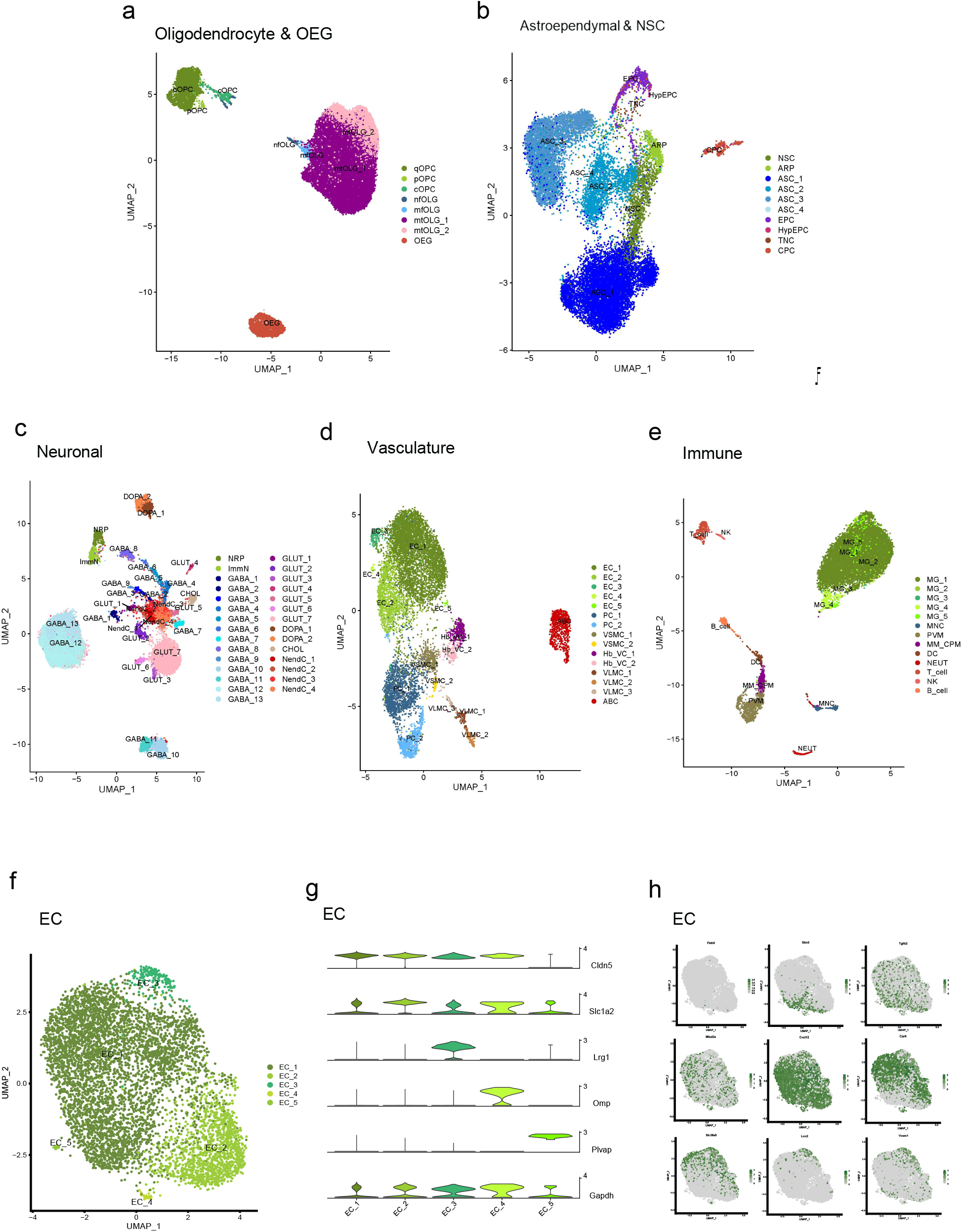
Characterization of cell types and sub populations. **a-e**. Subpopulation analysis of cell types grouped in 5 distinct cell classes: *Oligodendrocyte lineage & OEG, Astro-ependymal cells & NSC, Neuronal lineage, Vasculature cells*, and *Immune cells*. **f.** UMAP subpopulation analysis of EC clusters. **g.** Violin plot of delineating markers of ECs. **h.** UMAP overlay of EC zonation markers along the arteriovenous axis curated from the literature^2–4^. Markers in L-R order: *Fbln5, Gkn3, Tgfb2, Mfsd2a, Cxcl12, Car4, Slc38a6, Lcn2, Vcam1*.

As an example of this subclustering analysis, we identified 5 subpopulations of ECs in all animal types, thus revealing the extent of their heterogeneity. Specifically, we identified ECs only expressing classical markers such as *Cldn5* (EC_1), which represent the largest fraction of ECs, but also ECs positive for astrocytic markers, such as *Slc1a2* (EC_2)^46^, ECs expressing the mitogenic/neovascularization marker *Lrg1*^47^ (EC_3), ECs expressing the olfactory marker *Omp* (EC_4), and ECs specific to choroid plexus capillaries as denoted by the expression of *Plvap*^48^ (EC_5) (Fig 2f-g). As found in our previous study^42^ and by others^48,49^, our clustering analysis showed that there was no distinct separation of arterial, capillary, and venous ECs. However, select markers exhibited a zonation effect that further highlights the increased heterogeneity of ECs derived from different vascular beds. More specifically, as shown in Fig 2h, probabilistic programming of cell class assignment using arterial/capillary/venous markers characterized in recent studies ^48–50^ similarly displayed a clear zonation of ECs along the arteriovenous axis (Supplementary Fig 8).

### Heterochronic parabiosis reprograms the transcriptional landscape of multiple brain cell types

Parabiosis causes age-related phenotypic alterations in multiple distant organs and tissues, including the brain^51–53^. Specifically, adult tissues experience some degree of structural and functional improvement when exposed to young blood, and conversely, deterioration when exposed to old blood. Our prior work detailed transcriptional changes that take place in brain cells during aging^42^, but not all of these changes are necessarily of functional consequence. Therefore, we reasoned that heterochronic parabiosis provides a first-pass filter to help identify those aging-related genes whose expression are changes in the brains of parabionts are likely to be of functional consequence.

To explore the effects of heterochronic parabiosis with respect to brain aging at the transcriptional level, we performed differential gene expression (DGE) analysis. Specifically, we sought to identify the gene changes that are associated with the rejuvenation process in old heterochronic parabionts, and the gene changes that are associated with the aging acceleration process in young heterochronic parabionts. To accomplish this, we took into account the effects of the parabiosis surgery itself. We assessed DGE by leveraging the capability of a pseudobulk approach (see Methods) to allow for chained pairwise comparisons of parabiosis-induced rejuvenation (RJV) or aging acceleration (AGA) with respect to the parabiosis surgery and the physiological aging process^54–56^. This approach allowed us to create models beyond simple pairwise comparisons of heterochronic versus isochronic or heterochronic versus unpaired animals, followed by intersections of the paradigms and their set differences. However, for cross-referencing purposes, we also completed all relevant pairwise comparisons to directly compare the transcriptomes of any two animal types.

Overall, our chained pairwise model allowed us to make comparisons of heterochronic parabiosis gene signatures that either shifted more towards reversal or acceleration of aging. Consequently, two major DGE datasets were generated: (a) the RJV-associated and (b) the AGA-associated. More specifically, the RJV DGE dataset lists the old heterochronic genes (OY), taking into account the effects of the parabiosis procedure (OO) that are associated with a transition to a “more youthful” state (OX-YX). Conversely, the aging AGA DGE dataset lists the young heterochronic genes (YO) accounting for the effects of the parabiosis procedure (YY) and are associated with a transition to a “more aged” state (YX-OX).

Of the 20,900 total detected genes, 700 were significantly changed with aging in at least one cell type (false-discovery rate (FDR) ≤ 0.05), while 442 were significantly changed in RJV and 155 were significantly changed in AGA (Supplementary Tables 3-11). As expected, we did not capture significant transcriptional changes in all the identified cell types, potentially due to the low number of cells sequenced in some populations and the low levels of detected transcription and dropouts inherent to scRNA-seq analysis^57^. Nonetheless, our analysis also showed that the proportions of total DGEs per cluster that were common to aging and RJV varied from cell type to cell type, as did those that were common to aging and AGA (Supplementary Fig 9). The DGEs in common between RJV and aging and AGA and aging, along with genes unique to each paradigm, demonstrate that not all effects of aging are reversed or accelerated by parabiosis, and that heterochronic parabiosis may also work through genes and pathways independent of the aging process (Supplementary Tables 12-13).

Upon examination of the DGE datasets, a clear pattern of gene expression changes in RJV and AGA (FDR ≤ 0.05) was evident among our large cell populations (> 5000 cells) (Fig 3, Supplementary Tables 3-11). Specifically, OLG, ASC, GLUT, GABA, EC, and MG yielded the largest number of putative DGEs across the comparisons (Fig 3, Fig 4a, Supplementary Tables 3-11). While OLGs, which had the largest number of cells, had many aging and RJV DGES, they did not show many statistically significant changes in the AGA model. On the other hand, ECs, an average cluster in size, had the next largest number of gene changes in aging, RJV, and AGA genes followed by MG (Fig 4a). These data indicated that the EC population, most directly exposed to circulating factors, is highly affected in both the RJV and AGA paradigms. This is in accordance with our previous observations showing improvements in the brain vasculature after heterochronic parabiosis ^27^ and following GDF11 treatment ^29^ and further emphasizes the key role that ECs play in regulating communication between blood and brain^32^.

**Fig 3.**
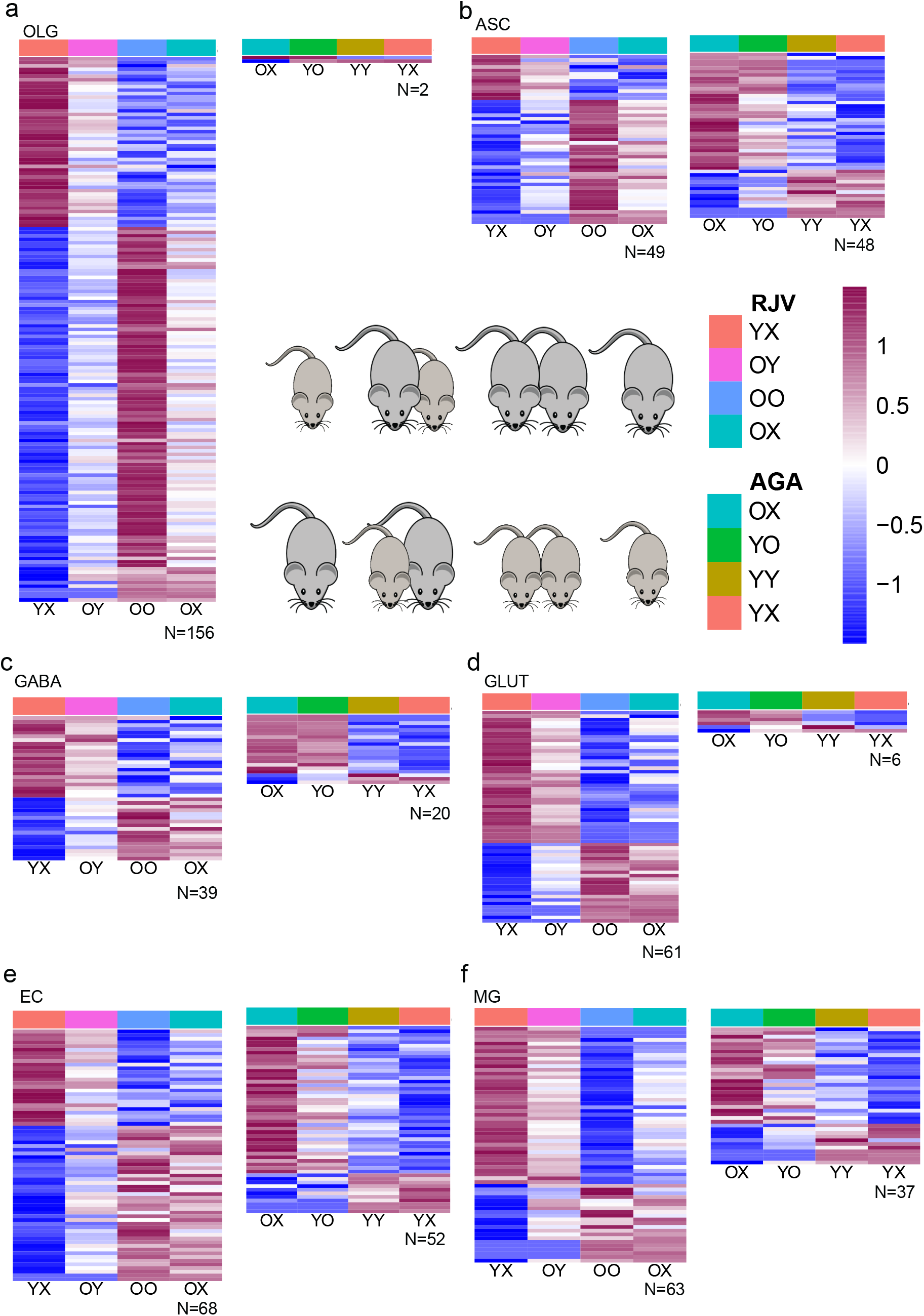
Differential gene expression (DGE) across major cell types revealed rejuvenation-associated (RJV) DGEs and aging acceleration-associated (AGA) genes. **a-f**. An FDR ≤ 0.05 was used to identify significant DGE genes. RJV framework depicts normalized gene expression changes across YX, OY, OO, OX. AGA framework depicts normalized gene expression changes across OX, YO, YY, YX. Genes are log2 z-scored scaled across rows and genes are ordered by logFC, with OLG (a), ASC (b), GABA (c), GLUT (d), EC (e), MG (f) in respective order. Color bar of the heatmap is reflects z-score, from negative (blue) to positive (magenta).

**Fig 4.**
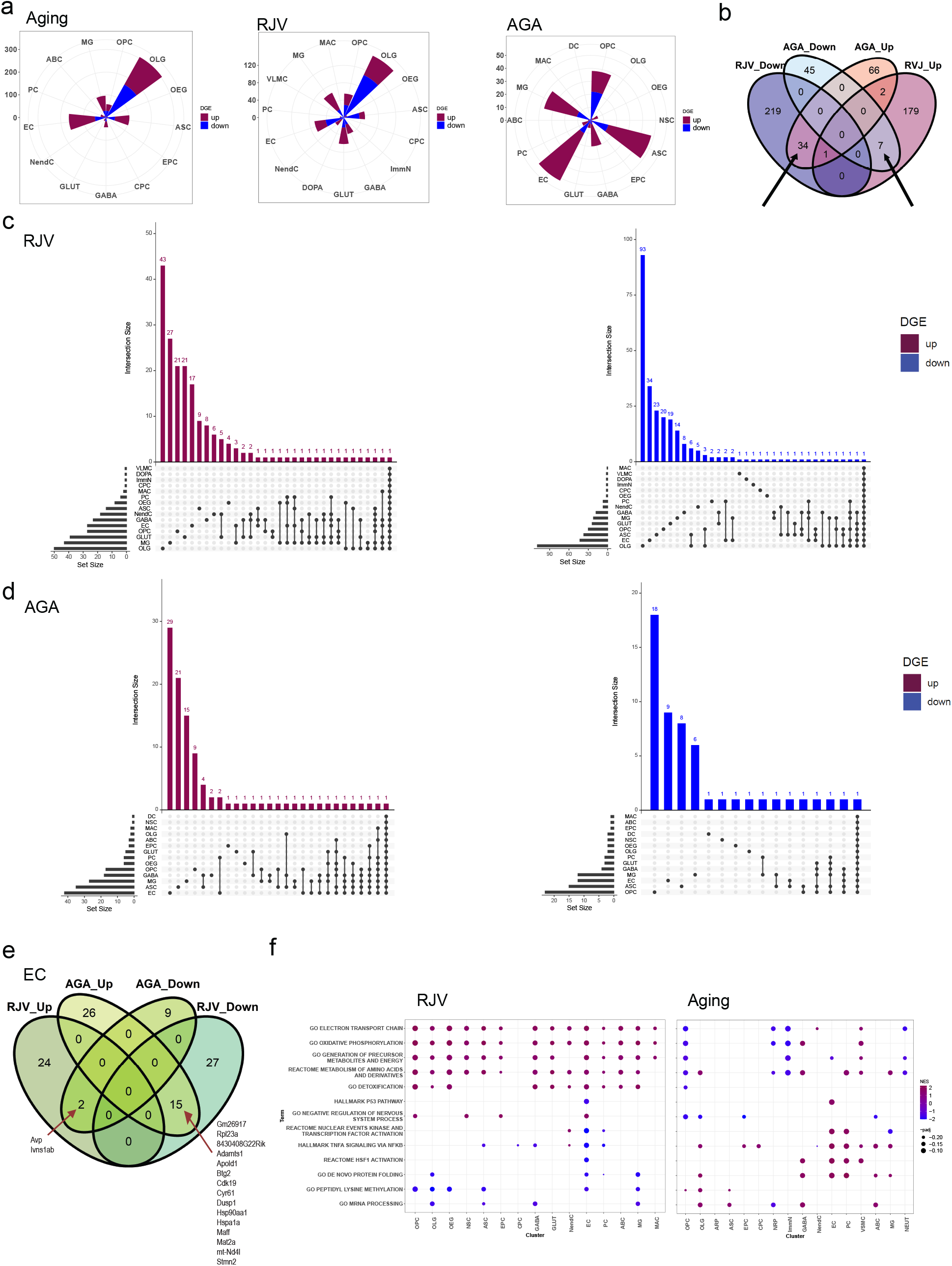
DGE characterization across cell types. **a:** Rose diagrams (circular histograms of number of DGEs) of Aging, RJV, AGA across all cell types at FDR ≤ 0.05, colored by direction of logFC (up magenta, down blue). **b.** Venn diagram of RJV and AGA DGEs across all cell types, demonstrating bidirectional logFC changes between the comparisons (depicted with arrows). **c-d.** Upset plot of FDR 0.05 DGE with positive logFC (upregulation) and negative logFC (downregulation) in both RJV and AGA. Top bar height reflects number of DGEs in the intersection, side bar width reflects magnitude of the set size. **e**. EC RJV and AGA DGEs Venn diagram split by logFC sign, revealing genes that reverse direction between comparisons. Arrows point to listed bi-directional genes. **f.** GSEA dot plots (padj ≤ 0.25) of representative terms across cell types in RJV and aging, with size of dot proportional to -padj, color by normalized enrichment score (NES) from negative (blue) to positive (magenta).

### Heterochronic parabiosis shifts gene signatures associated with both rejuvenation and aging acceleration

To find genes that are dysregulated across multiple cell types and to provide an overall perspective concerning transcriptional remodeling occurring after heterochronic parabiosis, we looked for key gene signatures that change in opposite directions in both RJV and AGA. Across all cell types, a total of 41 unique genes were found to change their expression levels bidirectionally in RJV and AGA, which corresponds to ~7% of all detected RJV and AGA DGEs. Specifically, 34 genes were downregulated in RJV and upregulated in AGA (Fig 4b), which corresponds to ~8% of all detected RJV DGEs, and ~22% of all detected AGA DGEs (Supplementary Table 14). To identify cell types that uniformly change the direction of expression of the same genes in the parabiosis-induced RJV process and/or in the parabiosis-induced AGA process, we created higher-dimensional Venn diagrams through upset plots (Fig 4c-d). This analysis identified certain pairs of cell types that shared multiple DGEs in the same direction in RJV (such as EC-MG and OLG-ASC), and in AGA (EC-PC and EC-MG) (Fig 4c-d, Supplementary Tables 15-16).

From all the analyzed cell types, ECs again demonstrated the greatest number of genes that change direction of expression (logFC) between RJV and AGA, further highlighting their susceptibility to the aging process and their potential for manipulation. For example, the transcription factor (TF) *Maff*, which is highly upregulated with aging in ECs, was found downregulated in RJV and upregulated in AGA (Fig 4e, Supplementary Table 14). Similarly, the aging-upregulated genes *Hsp90aa1* and *Hspa1a*, which encode heat shock response proteins, *Adamsts1*, which is induced upon sheer stress^58^, *Apold1*, which is responsive to hypoxia/ischemia^59^ and LPS treatment^32^, *Cyr61*, which encodes an ECM protein involved in angiogenesis^60^, *Dusp1*, which participates in EC migration^61^ along with *Stmn2*, a gene involved in the microtubule organization, were all downregulated in RJV and upregulated in AGA (Fig 4e, Supplementary Table 14). In the opposite direction, the aging-downregulated gene *Avp*, the expression of which is known to decrease in neurodegeneration^62^ was upregulated in RJV and downregulated in AGA. Similarly, *Ivns1ab*, known to stabilize the cytoskeleton and protect from cell death induction due to actin destabilization ^63^, followed the same trend, suggesting a structural improvement of ECs upon heterochronic parabiosis (Fig 4e, Supplementary Table 14).

The DGE analysis also showed that the transcription factor *Klf6*, which was one of the most upregulated genes with aging in ECs, was differentially downregulated in RJV, as was its downstream target *Smad7*^64^ (Supplementary Tables 3-4). Considering that *Klf6* expression is known to be regulated by vascular injury^65,66^, we further characterized its transcriptional changes by RNA *in situ* hybridization. As shown in Fig 5a-b, *Klf6* expression was indeed found to be highly upregulated with aging in ECs (*Pecam1^+^*). Heterochronic parabiosis reversed this change in the old parabionts, bringing its transcriptional levels of Klf6 close to those seen in young ECs (Fig 5b).

**Fig 5.**
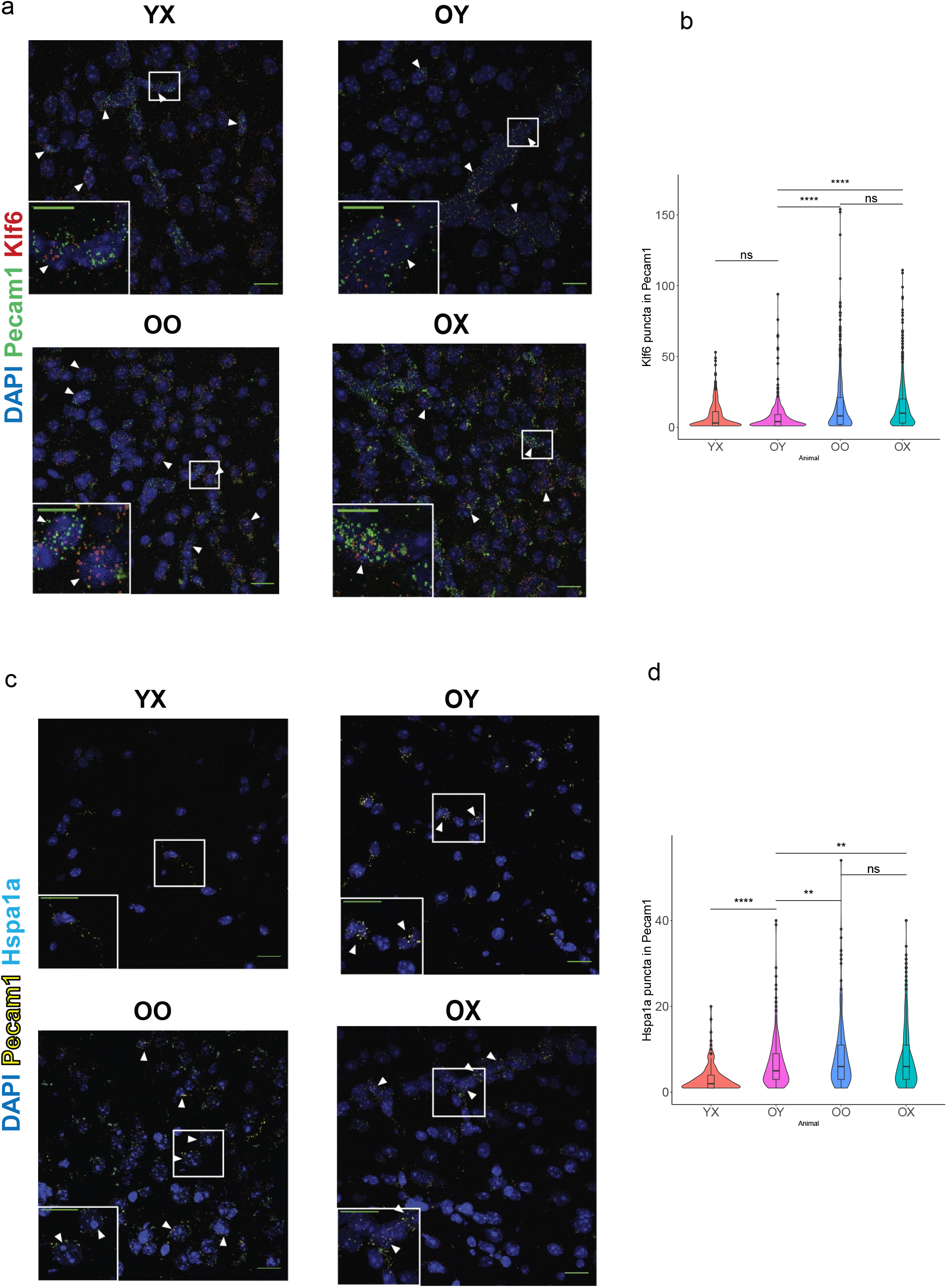
RNA *in situ* hybridization assays showing Aging and RJV reversal of key aging-associated genes. **a.** Representative RNA images of mouse cortices showing *Klf6* puncta in *Pecam1*^+^ ECs in YX, OY, OO, OX mice. Scale bars represent 50μm for each subplot, with 10μm inset. **b.** Violin plot representation of RNA quantitation by two-tailed Welch’s t-test for significance. P-values for OY-YX: 0.473, OY-OO: 5.328e-19, OY-OX: 4.900e-28, OO-OX: 0.349. (ns p-value > 0.05, * p-value ≤ 0.05, ** p-value ≤ 0.01, *** p-value ≤ 0.001). **c.** Representative RNA images of mouse cortices showing *Hspa1a* puncta in *Pecam1*^+^ ECs in YX, OY, OO, OX mice. Scale bars represent 50μm for each subplot, with 10μm inset. **d**. Violin plot representation of RNA quantitation by two-tailed Welch’s t-test for significance. P-values for OY-YX: 3.535e-16; OY-OO 0.008, OY-OX: 0.006, OO-OX: 0.686. (ns p-value > 0.05, * p-value ≤ 0.05, ** p-value ≤ 0.01, *** p-value ≤ 0.001).

Taken together, our computational analysis identified numerous aging-related genes across multiple cell types including ECs whose expression were restored in old mice and/or disrupted in young mice following heterochronic parabiosis, suggesting their potential involvement in the rejuvenation and/or aging acceleration process, respectively. These data are again consistent with the notion that parabiosis is likely to act in part by regulating processes important to vascular structure and health.

### Heterochronic parabiosis reverses aging-induced pathways

To profile transcriptional changes beyond genes identified by DGE analysis, we performed an implementation of Gene Set Enrichment Analysis (GSEA)^67^ to reveal biological processes and molecular pathways associated with aging, RJV and AGA. Composite ranks were calculated for each gene based on FDR and logFC. As with our previous study on old and young animals^42^, the new ranking metric used yielded a large number of significant terms for most cell types (Supplementary Table 17). Specifically, in the RJV model, the most gene sets were observed in OPC, OLG, OEG, ASC, GABA, GLUT, NendC, EC, and MG, whereas in the AGA model, only OPC, OLG, and PC yielded significant terms. This further reinforced our observation that heterochronic parabiosis induces stronger aging-related gene signature changes in the old parabionts.

Among the identified changes, GSEA revealed key ontologies that were downregulated in aging and upregulated in RJV across multiple cell types. For example, pathways related to mitochondrial activity, such as *GO Oxidative Phosphorylation*, and *GO Electron Transport Chain*, as well as oxidative stress homeostasis and metabolism pathways, such as *GO Detoxification* (a category that includes genes for transporters known to function at the blood-brain barrier)*, GO Generation of Precursor Metabolites and Energy*, and *Reactome Metabolism of Amino Acids and Derivatives* followed this trend (Fig 4f, Supplementary Table 17). RJV downregulated pathways like *GO DNA Conformation Change* and *GO Peptidyl Lysine Methylation*, and further demonstrated that the epigenetic machinery is functionally perturbed in various cell populations with parabiosis (Fig 4f, Supplementary Table 17).

In addition to the above ontologies, ECs displayed a clear pattern of normalized enrichment score (NES) sign reversal between aging and RJV, corresponding to our DGE profiling. Changes in mitochondrial and metabolic pathways were also upregulated in RJV. Processes that were downregulated in ECs include inflammatory pathways like *Hallmark TNFA Signaling via NFKB*, and apoptosis and senescence-associated *Hallmark P53 Pathway*. Likewise, the protein homeostasis (proteostasis) associated pathways, such as *Reactome HSF1 Activation* and *GO De Novo Protein Folding*, were found to be upregulated with aging and downregulated in RJV (Figure 4f, Supplementary Table 17). Collectively, these data suggested that heterochronic parabiosis changes the metabolic profile, improves proteostatic machinery, and reduces aging-associated apoptosis or senescence to improve EC function, consistent with recent findings in aortic ECs^68^.

We then focused on further exploring the decline of proteostasis, an acknowledged hallmark of aging, in ECs. Collapse of complex proteostasis networks in aging results in the formation of misfolded proteins and the accumulation of protein aggregates^69^. Our pathway analysis revealed upregulation of several stress-inducible pathways in ECs that are presumably activated in response to misfolded protein accumulation in aging and suppressed in RJV (Fig 4f, Supplementary Table 17). To explore this, we examined the gene expression levels of *Hspa1a*, which encodes a stress-inducible heat shock protein. At the DGE level, *Hspa1a* was found to be upregulated in aging and downregulated in RJV in EC (Supplementary Tables 3-4). To verify this change, and to further demonstrate that defects in proteostasis are mitigated by heterochronic parabiosis, we performed RNA *in situ* hybridization. As shown in Fig 5c-d, we detected a significant decline in the number *Hspa1a* puncta expressed by *Pecam1*^+^ cells between old heterochronic parabionts and both aged mice and control old isochronic parabionts.

### Heterochronic parabiosis activates global remodeling of gene regulatory networks

In an attempt to identify transcription factors (TFs) that regulate the response in heterochronic parabiosis, regardless of whether their own gene expression changes during aging, RJV or AGA as identified by DGE analysis, we utilized the SCENIC approach. This framework allowed us to detect transcriptionally-active gene regulatory networks (GRNs), which are comprised of TFs and their downstream effector genes^70^. Each animal type was profiled individually to identify putatively active GRNs, stratified by cell class. The young unpaired and young isochronic animals had fewer cell types with higher frequencies (>100) of higher (>1) GRN activity scores than young heterochronic animals (Fig 6a). VLMC, a newly identified perivascular-like cell type of the brain vasculature^46^, frequently had high activity across all animal types (Fig 6a). In old heterochronic animals, VLMC, NEUT, ASC, and DOPA had higher frequencies of high (>1) GRN activity scores than old unpaired animals (Fig 6a, Supplementary Table 18). These putative changes in GRN activity may suggest an increase in global remodeling in the old and young heterochronic mice to reflect RJV and AGA, respectively.

**Fig 6.**
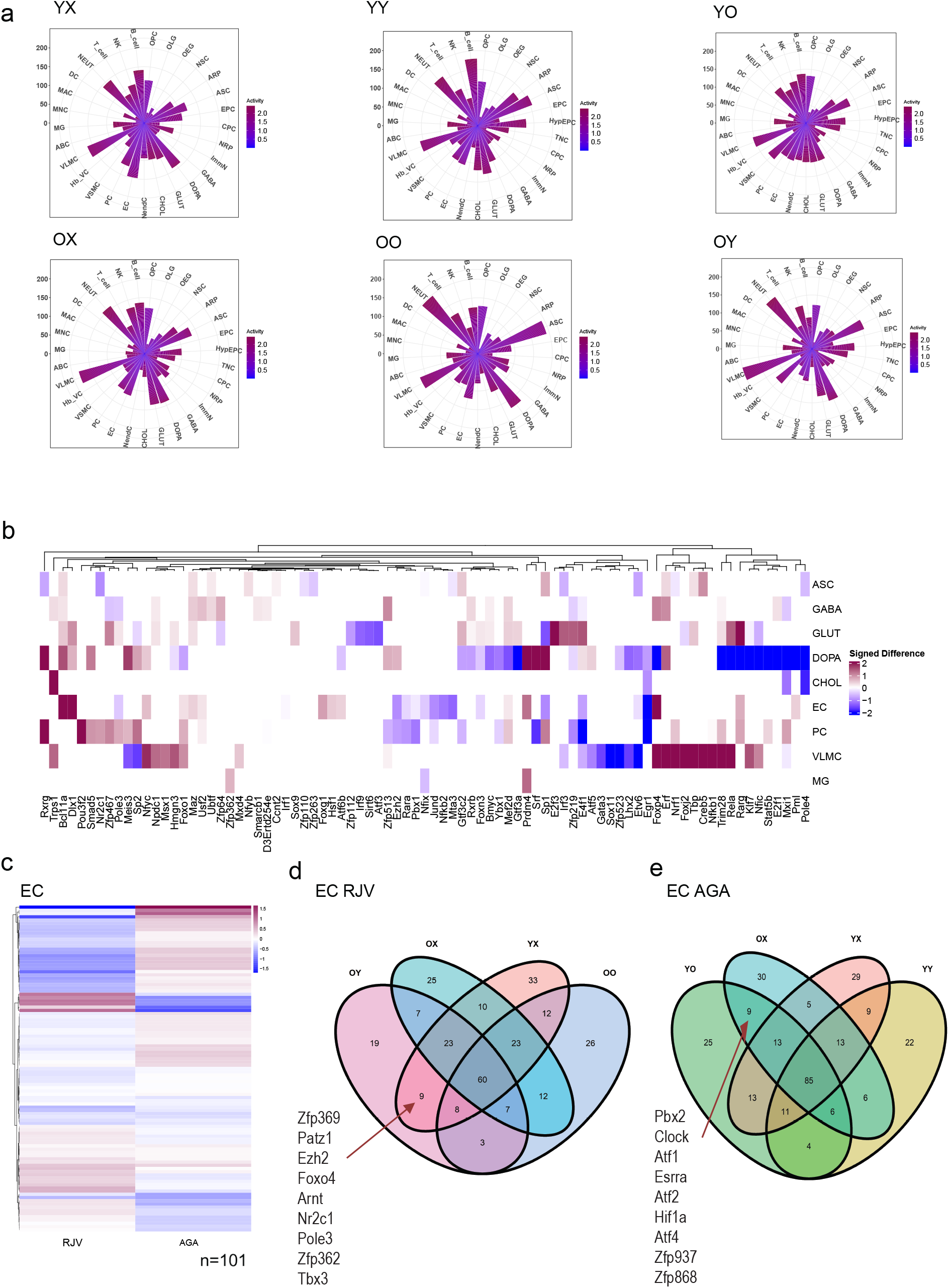
Gene regulatory analysis reveals transcriptionally active cell types and gene regulatory network (GRN) reversals between RJV and AGA. **a:** Rose diagrams of GRN scores per cell type across YX, YY, YO, OX, OO, OY. Color scale from blue to magenta reflects the degree of GRN activity (see Methods for details). **b**. Difference heatmap of active GRN transcription factors (TFs) corresponding to RJV/AGA logFC change sign. Magnitude is the absolute magnitude of the difference, direction is positive for upregulation in RJV (magenta), negative for upregulation in AGA (blue). **c**. Heatmap of EC active GRN TFs plotted by RJV and AGA logFC, with upregulation magenta, downregulation blue, clustered with Euclidean distance, average linkage. **d-e:** Venn diagrams of EC RJV and AGA animal frameworks’ active GRN TFs. Arrows point to those TFs in common between OY and YX and YO and OX respectively.

RJV via parabiosis has a distinct transcriptional landscape that follows the opposite direction of expression in AGA. The signed absolute difference in logFC identifies TFs that are bidirectional between RJV and AGA, meaning that they have increased expression in RJV and downregulation in AGA, or vice versa. DOPA demonstrated the most extreme instances of TFs that change direction, followed by VLMCs, then ECs (Fig 6b, Supplementary Table 14). Considering that ECs displayed many bi-directional TFs, we further explored their entire transcriptional landscape. Of the EC TFs in old and young heterochronic brains, the majority demonstrated opposite regulation in RJV and AGA (Fig 6c). TFs in common between old heterochronic and young unpaired ECs that reflect the RJV construct included essential regulators of EC function such as *Tbx3*^71,72^, *Foxo4*^73^ which is a member of the FOXO family of TFs that are components of a fundamental aging regulatory pathway, *Patz1*, which has been implicated in regulating p53 levels and senescence in ECs^74^, and *Arnt*, which participates in aryl hydrocarbon receptor signaling and is involved in several aspects of vascular biology^75^ (Fig 6d). Likewise, young heterochronic and old unpaired ECs that reflect the AGA construct shared the hypoxia response genes *Atf1, Atf2*, and *Hif1a*, indicating a stressed EC profile, as well as *Atf4* which has been involved in angiogenesis^76^ (Fig 6e).

### Heterochronic parabiosis alters aging-associated intercellular communication networks and interactions

Altered intercellular communication is one of the hallmarks of aging^1^. While many studies have examined the actions of blood-borne factors on CNS cells, few have looked at factors secreted by the CNS cells themselves and how they are modified by aging. The importance of such secreted factors has been clearly documented, where dysregulated intercellular and inter-tissue signaling exacerbates inflammation and degeneration^77–80^.

To delineate the potential effects of heterochronic parabiosis on intercellular communication within the brain, we built networks of putative ligand-receptor interactions across all the identified brain cells for each animal type. The tool CellChat^81^ quantifies ligand-receptor interactions based on the law of mass action, wherein the probability of a ligand and receptor signaling is modelled by their average expression values as well as their cofactors. Use of CellChat allows construction of a theoretical framework of nearest neighbors where the factor released by each cell type and the distance to its putative receptor are taken into account^81^.

We first measured the total number of interactions for each animal type to elucidate the number of cell-cell communication connections (Fig 7a, Supplementary Table 19). We found that the old heterochronic parabionts exhibited fewer putative connections than the old unpaired and old isochronic animals, while the young heterochronic parabionts exhibited more putative connections than in the young unpaired and young isochronic animals. To disentangle the complexity of these networks, and to reveal the intercellular interactions that could potentially be associated with rejuvenation and aging processes, we examined those ligand-receptor pairs that change with both aging and heterochronic parabiosis. We focused on 201 cell-cell communication pairs found only in old heterochronic and young unpaired brains, and not in old isochronic and old unpaired brains, which reflects the RJV paradigm. Similarly, we identified 392 interactions shared between young heterochronic and old unpaired brains, but not found in young isochronic and young unpaired brains, to reflect the AGA paradigm (Fig 7b).

**Fig 7.**
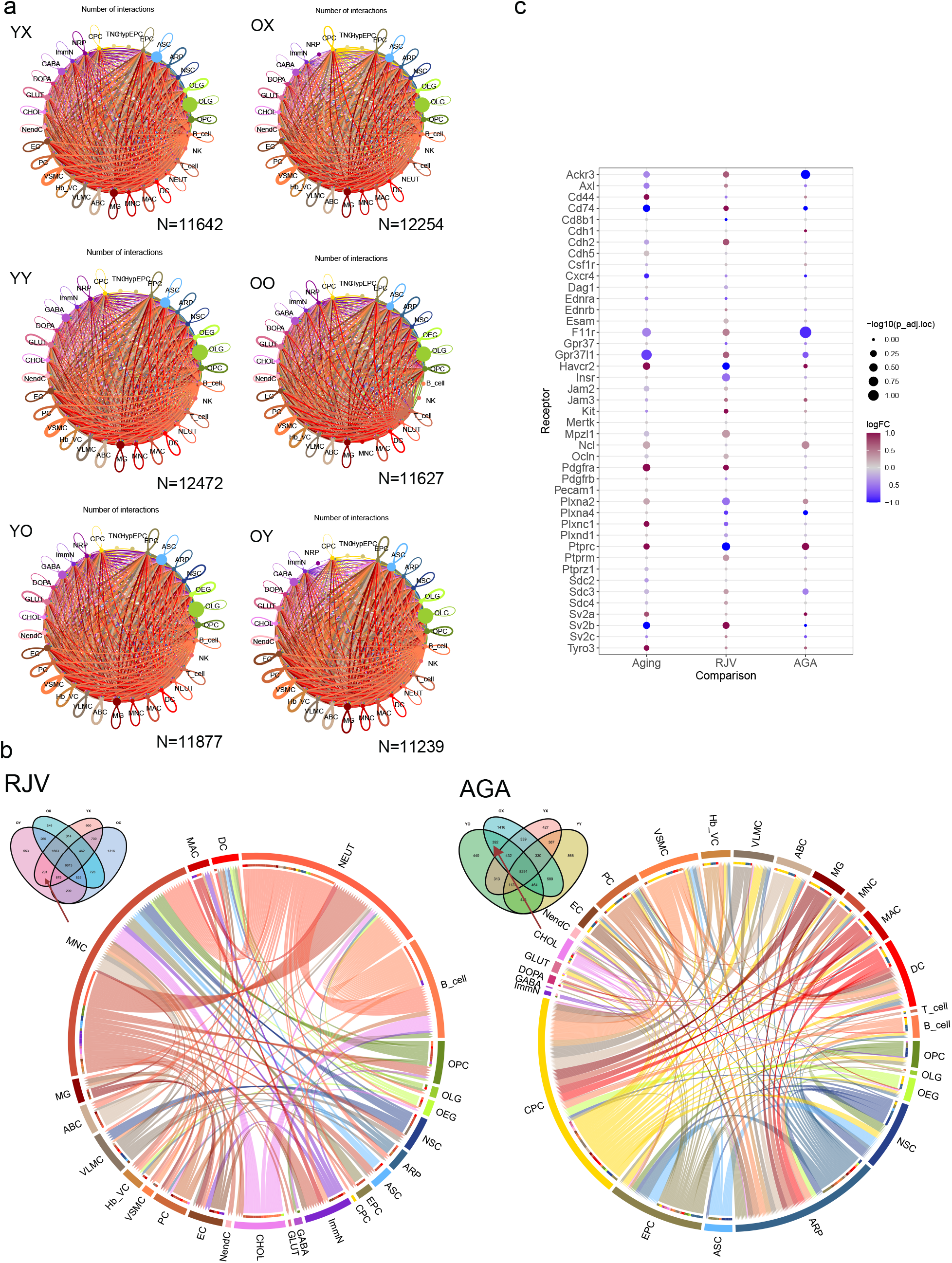
Cell-cell communication is affected with aging and parabiosis. **a.** Summarization network graphs of the number of ligand-receptor interactions between cell types in YX, YY, YO, OX, OO, and OY mice. Node size is proportional to cell population size. Edge width and transparency of color are proportional to the number of all edges between a set of nodes. **b**. Chord diagrams representing the informatically-predicted unique source:target:receptor:ligand pairings identified only in the rejuvenation model OY and YX (Venn diagram inset, left panel) or in the aging acceleration model of YO and OX (Venn diagram inset, right panel). **c**. For all identified EC receptors, DGE metrics are shown for the aging, RJV, and AGA paradigms. Node size is inversely proportional to the adjusted p-value, and node color is scaled by intensity of logFC from blue (negative, downregulation) to magenta (positive, upregulation).

In RJV, we observed more signaling among various blood cell types, suggesting their overall contribution to the rejuvenation process (Fig 7b). This may also be due to the fact that we did not perfuse the mice prior to brain tissue collection because of technical challenges with the parabiosis surgery. However, in RJV, we also discovered connections triggered by VLMC and ABC, both of which act physiologically as barrier cells. For example, we found that VLMC and ABC potentially signaled to ASC, EC, and MG via a variety of factors including *Mdk, Ptn*, and *Wnt5a*, while they received signals from NSC, NRP, and VSMC mediated for example by *Notch* ligands (Fig 7b, Supplementary Table 19). Conversely, in AGA, more signaling was triggered by CPC. Specifically, CPC signaled to OPC and various neuronal cell types (DOPA, GLUT, CHOL) via *Nrxn2*, while they received signals from VSMC and GLUT. In this paradigm, we also observed more signaling triggered by EPCs, which are cells that also act physiologically as barriers as they form the epithelial lining of the ventricles (Fig 7b, Supplementary Table 19). Taken together, this computational analysis highlighted the prominent roles for various barrier cells in the brain parenchyma, as well as CPCs, in processes accompanying or mediating the effects of parabiosis.

Following our previously published approach^42^, we pursued ligand-receptor interactions determined by encapsulating FDR and logFC as the metric for putative association. This computational analysis showed that there are certain aging-induced changes in cell-cell communication, including those triggering excessive proinflammatory responses, that were reversed in the old heterochronic parabionts and/or potentiated in the young heterochronic parabionts. As an example, we detailed interactions involving ECs, which are known to secrete signals that regulate neurogenesis^82^ and recently were reported to be implicated in signaling networks with OLG lineage cells^83^. When the interactions among EC ligands and their putative cognate receptors in OLG and NRP were explored, we delineated a clear pattern of connection direction reversal between aging and RJV for most ligands, as well as between RJV and AGA, with AGA exhibiting a direction similar to aging (Supplementary Fig 10). We identified several secreted factors that could mediate these intercellular relationships. Of those, cytokines/inflammatory mediators such as IL4^84^, IL18^85^, CXCL12^86^, and TNFa^87^ are known to affect myelination. Growth factors BDNF^88^, TGFb^89^, and BMP4^90^ are also shown to have strong effects on myelination. With respect to factors that might effect neurogenesis, we identified CXCL12^91^ and TNFa^92^ as ligands from ECs. We also found that ECs are a potential source of BDNF, which is well-known as a significant regulator of neurogenesis^93^, ANGPT2^94^, and GDF11, which has also been recently demonstrated by our lab to regulate neurogenesis^95^.

Another way of investigating the data is to look for aging/RJV-dependent changes in the levels of cellular receptors. We applied this analysis to ECs, known to be highly influenced by factors found in blood and recognized to be subject to interactions with PC and ASC. The genes encoding receptors found in the cell-cell communication analysis are shown in Fig 7c, with their DGE adjusted p-values and logFC extracted for the aging, RJV, and AGA paradigms. The receptors include membrane proteins that constitute parts of junctional complexes, such as JAM protein that respond to sheer stress, mediate vascular remodeling, and regulate EC barrier function^96^. We also identified a variety of receptors known to participate in angiogenesis, including TAM receptors, which mediate GAS6 signaling^97,98^, syndecans, which are co-receptors for VEGF and other ligands^99,100^, plexins, which regulate angiogenesis by binding to semaphorins^101,102^, CXCR4, a receptor for CXCL12 which affects EC proliferation^103^ and leukocyte extravasation^104^, and CD44, a receptor for ECM proteins^105,106^.

Collectively, our computational analysis identified numerous cell-cell communication networks that are perturbed during the aging process and are modified upon heterochronic parabiosis. This may point to sets of intercellular interactions that can be functionally manipulated for targeted aging intervention. Our computational analyses also highlighted the significance of ECs as a potential target for therapeutics, since its intercellular interactions are affected by both aging and heterochronic parabiosis.

### Heterochronic parabiosis regulates the senescence state

The effects of parabiosis on the aging process with respect to cellular senescence, a well-known and widespread hallmark of aging^1^, are beginning to be recognized^37,107^. To explore this in more detail, we performed an implementation of GSEA analysis with a reference gene set of literature-defined senescence-associated genes (Supplementary Table 20)^107,108^ against the pre-ranked aging, RJV and AGA gene sets to determine functional enrichment of senescence in these paradigms, and their directions.

The literature-curated gene set was able to discern patterns of upregulation and downregulation of senescence-associated genes in aging, RJV, and AGA. Across 20 of the 29 examined cell populations, we observed directionality reversal between aging and RJV based on NES (Fig 8a, Supplementary Table 21). In the vast majority of cell types (OPC, OLG, ARP, ASC, EPC, CPC, GABA, DOPA, EC, PC, MG, MNC, NEUT, B-cell), aging had a positive enrichment score, indicating upregulation of senescence consistent with prior studies^109^, whereas RJV had a negative enrichment score indicating reduced levels of senescence. The NES direction was the same in aging and AGA for 20 cell types, indicative of recapitulation of the aging process in AGA (Fig 8a). However, beyond computed gene set directionality as measured by NES, many clusters were not statistically significant for senescence enrichment at an FDR ≤ 0.25, but we captured 11 significant cell types in aging, 7 in RJV, and 10 in AGA (Fig 8a, Supplementary Table 21).

**Fig 8.**
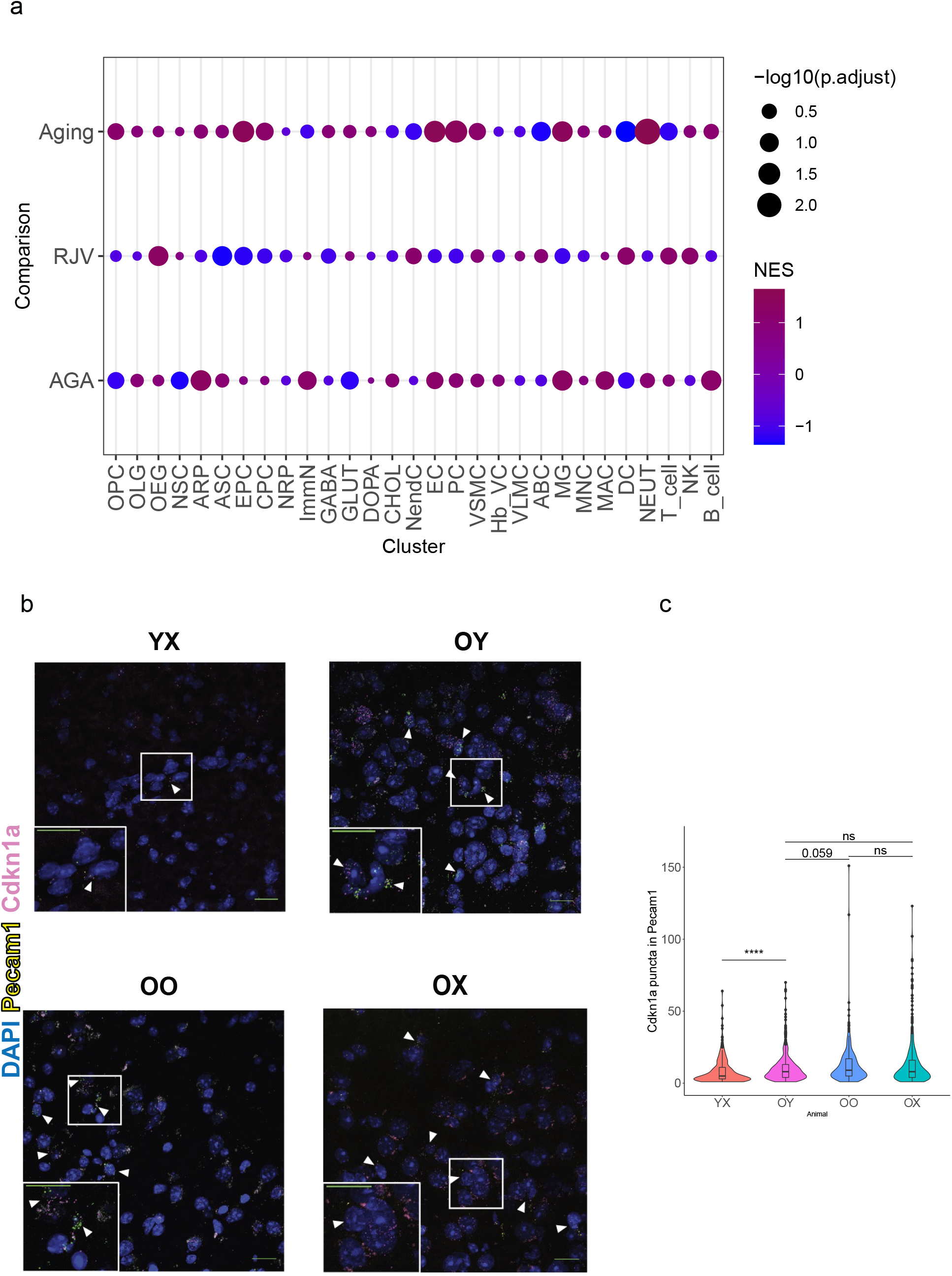
Senescence status demonstrated shifts in aging and RJV. Dot plot representation of senescence-associated marker genes curated from the literature^5 6^, permuted against each cell type in Aging, RJV, and AGA with fGSEA. The inverse log10 adjusted p-values (Benjamini-Hochberg) reflect the size of the dot, and Normalized Enrichment Scores (NES) reflect color from blue (negative enrichment) to magenta (positive enrichment). **b.** Representative RNA *in situ* images of mouse cortices showing *Cdkn1a* puncta in *Pecam1*^+^ ECs in YX, OY, OO, and OX mice. Scale bars represent 50μm for each subplot, with 10μm inset. **c.** Violin plot representation of RNA *in situ* data quantitation. Wilcoxon rank sum exact test for significance. P-values for OY-YX: 4.374e-06, OY-OO 0.059, OY-OX 0.581, OO-OX: 0.153. (ns p-value > 0.05, * p-value ≤ 0.05, ** p-value ≤ 0.01, *** p-value ≤ 0.001).

To further examine whether the senescence-associated gene signature is indeed modified by parabiosis, we performed RNA *in situ* hybridization. Specifically, we evaluated the gene expression changes in the regulation of the senescence-associated gene *Cdkn1a* in ECs. We observed a decrease of *Cdkn1a* expression in ECs (*Pecam1^+^*) in the old heterochronic parabiotic brains compared to the old isochronic brains (Fig 8b-c).

Taken together, our computational analysis indicates that heterochronic parabiosis modulates cellular senescence-associated genes in multiple brain cell types and suggests the possibility that the cellular senescence regulation itself may contribute, at least partly, to the positive and negative effects of parabiosis in the brains of old and young parabionts respectively.

## Discussion

Over the past few years, studies replete with compelling evidence have demonstrated that aging is a highly dynamic and malleable process, with several types of treatment reported to rejuvenate tissues and organs and in turn, extend organismal healthspan and lifespan^110,111^. Among these interventions, heterochronic parabiosis appears to be one of the most effective. Despite this, the mechanism(s) by which parabiosis acts at the cellular and molecular levels to improve tissue function remains elusive.

Here, we present a comprehensive single-cell survey of the gene expression changes that occur in the aged and young mouse brain following heterochronic parabiosis. Having previously described, also at the single cell level, the changes in gene expression that accompany aging ^42^, our main goal was to understand how gene expression is modified after old and young brains are exposed to complex sets of circulating factors associated with heterochronic parabiosis. In principle, this allows us to understand which cell types and which genes are associated with functional improvement or deterioration of the CNS.

We assessed the cellular complexity of the parabiotic brains and showed that cell identity and composition were largely retained. Specifically, we found the same cell populations across all parabiotic brains, without detection of unique cell clusters associated with certain animal types. We also observed that the number of cells in each type and subtype did not significantly change with heterochronic parabiosis. Thus, parabiosis-induced phenotypic and functional changes in the brain are not mainly due to shifts in the number of certain cell types, but rather due to changes in their expression profiles. However, as tissue dissociation is an inherent part of the single cell sequencing workflow, this aspect might have resulted in non-uniform sampling problems^112^. Therefore, we cannot completely rule out the possibility that parabiosis does affect the number of certain cell populations, as has been previously described^24,27^. Future studies employing single-nuclei sequencing approaches^113^ and spatially-resolved transcriptomics^114^ may shed more light on this matter.

We explored the gene expression changes associated with aging and heterochronic parabiosis in all identified brain cell types. Due to the high complexity of these datasets, we utilized a chained pairwise comparison framework that allowed us to effectively reveal which aging-induced changes were modified by heterochronic parabiosis in both the rejuvenation and aging acceleration paradigms. We analyzed those genes and pathways that modified the old heterochronic brains exposed to young blood to resemble their more youthful counterparts, and those that modified the young heterochronic brains exposed to old blood to become more similar to the aged brain. To this end, we identified the primary cell types exhibiting these changes and revealed the common and cell-type-specific aging-induced signatures and transcriptional programs that were rescued following parabiosis in the old brains and/or disrupted in the young brains. As corroborated by recent reports, our data provide evidence that heterochronic parabiosis effectively modulates multiple manifestations of canonical aging hallmarks, including altered intercellular communication^79^, loss of proteostasis^115^, defects in mitochondrial dysfunction^68,116–118^ and cellular senescence^37,107,119,120^.

Our analyses suggest that the modulation of the aging process is mediated by reprogramming of the associated transcriptional signatures across multiple brain cell types. In support of this notion, aging-induced changes in the epigenetic status of the aged mouse brain^121^ as well as of the aged liver and blood tissues^122^ were recently reported to be reversed following heterochronic parabiosis. The reprogramming of the transcriptome presented here is somewhat similar to the reprogramming recently reported to take place in rats subjected to caloric restriction^123^. These findings raise the exciting possibility that both interventions could promote tissue rejuvenation by mitigating the appearance of similar aging-associated epigenetic alterations, and consequently their induced transcriptional changes. Thus, future studies investigating the exact epigenetic regulators and mechanisms that are responsible for these types of changes will be of high importance^111,124^.

Our data also support the view that parabiosis exerts effects independent of the aging process, an aspect incorporated into our analysis. As some of these effects are likely to have negative consequences^125^, it can be argued that the positive effects of young blood factors exposure can overcome not only aging-driven changes in gene expression, but also processes such as systemic inflammation and stress stimulated by the parabiosis surgery itself^16,126^. Moreover, parabiosis-driven effects may be mediated through means other than reversing the effects of normal aging. In recognition of this possibility, we designate the DGEs from the chained pairwise comparisons that have no overlap with aging DGEs (Supplementary Tables 12-13).

One of the primary discoveries of our study is that the EC transcriptome is dynamically regulated by both aging and heterochronic parabiosis. We found that ECs, when compared to other brain cell types, exhibited the highest fraction of aging-related genes that were rescued following heterochronic parabiosis in the old brain, and similarly, the highest fraction of aging-related genes that were disrupted following heterochronic parabiosis in the young brain. This finding supports our previous research that the vasculature is strongly affected by aging and disease and is capable of regrowth after heterochronic parabiosis^27^ or systemic GDF11 treatment^29^. We observed that a subset of ECs was classified as mitogenic, expressing high levels of *Lrg1* and *Lcn2* (Fig 2f-g) and it is reasonable to speculate that the growth of these cells, whose growth is probably prevented or suspended by the inflammatory environment of the aged brain, may be among the cells which respond to these interventions. In addition, while proteostasis in brain ECs has not been thoroughly investigated, they are apparently long-lived cells^32,127^ and, like neurons, might therefore accumulate protein aggregates with age^128^, potentially compromising their function. Finally, as previously shown, ECs become senescent with age^129,130^, but parabiosis may reverse that phenotype as well. Taken together, these findings underline the strong susceptibility and malleability of ECs, which are directly exposed to all secreted factors derived from both brain parenchyma and blood, to adapt to changes in their microenvironment, which is consistent with prior observations from our lab^27,29^ and others^32,50,68,79^. Therefore, ECs, despite comprising less than 5% of the total number of brain cells^131^, are a promising and accessible target for the treatment of aging and its associated diseases^132^.

Collectively, our gene expression matrices provide unprecedented data that can be further explored to form working hypotheses for follow-up studies. Similar to other recent reports^118,123,133^, our study advances fundamental understanding of the mechanisms underlying the aging process and potential interventions that go beyond descriptive studies of cell states. Our work extends current knowledge regarding the effects of heterochronic parabiosis on the aging process and supports the key role of the brain vasculature in mediating these effects. Effectively, in one, albeit complex procedure, parabiosis improves many of the individual processes, such as mitochondrial dysfunction, proteostasis collapse, and cellular senescence, that are usually targeted separately by therapeutic interventions. Altogether, our study may provide a path towards identifying new single therapeutics avenues with multiple beneficial effects.

## Data Accessibility

Data from this study are currently hosted on a web portal at: https://hmsrsc.aws.hms.harvard.edu/parabiosis

## Acknowledgements

We would like to acknowledge Tomotaka Okino and his team at Ono Pharmaceuticals for fruitful discussions during the progress of this work. We are grateful to K. Pritchett-Corning, F. Rapino, K. Pfaff, I.A. Gampierakis, M.H.C Florido, S. Ghosh, and J. LaLonde for their advice and help in different aspects of the study. We also thank the staff members of the Harvard Biolabs Animal Facility, the Harvard Center for Biological Imaging, and the Harvard Stem Cell and Regenerative Biology Histology Core for their continuous support and assistance. Harvard Medical School Center for Computational Biomedicine and Research Computing host the web portal. This work was supported by Ono Pharmaceutical Co., Ltd, the Stanley Center for Psychiatric Research, the Klarman Cell Observatory of the Broad Institute of MIT and Harvard, Howard Hughes Medical Institute, an award from the Glenn Foundation, NIH/NINDS grant 1R01NS117407, NIH/NIA grant 5R01AG072086, NIH grant T32 DK007529, and the Simons Foundation. The funders had no role in the study design, experiments performed, data collection, data analysis and interpretation, or preparation of the manuscript.

## Author Contributions

M.X. and L.L.R. conceived the study; M.X., K.M.H. and L.L.R. designed the study; M.X. performed the parabiosis experiments; M.X., X.A., D.D., and L.N. performed the single cell RNA-seq experiments; K.M.H., S.L.L. and J.Z.L. processed the single cell RNA-seq data; K.M.H. developed the computational framework and performed all associated analyses; M.X. and K.M.H. interpreted the data; R.M.G., C.O., M.S.; S.S., and C.G. designed and/or performed validation experiments;; J.M.G performed the blood-chimerism experiments and analysis; K.M.S. assisted in the development of transcriptional networks; K.M.H. built the online portal. M.X., K.M.H., A.J.W., A.R., and J.Z.L. supervised aspects of the study; L.L.R. directed the study; J.Z.L. and L.L.R. secured funding; M.X. and K.M.H. wrote the original draft of the manuscript; C.O., S.K.S, S.M.B., J.Z.L., A.R., and L.L.R. provided critical feedback and/or edited the manuscript; All authors reviewed the manuscript and approved its submission.

## Methods

### Animals

C57BL/6J inbred male mice (JAX no. 000664; CD45.1^-^CD45.2^+^) and B6.SJL-Ptprc^a^ Pepc^b^/BoyJ male mice (JAX no. 002014; CD45.1^+^CD45.2^-^) were housed in the Harvard Biolabs Animal Facility under standard conditions. All experimental procedures were approved in advance by the Institutional Animal Care and Use Committee of Harvard University (AEP no. 0-23) and are in compliance with federal and state laws. On the day of sacrifice, young mice were 3-4 months (13-15 weeks) of age, and old mice were 20-22 months (80-87 weeks) of age.

### Parabiosis

Parabiosis surgeries were performed as previously described^27,134^ with a few modifications. In brief, mice were sedated with controlled isoflurane anesthesia and placed on heating pads to prevent hypothermia. Ophthalmic ointment was applied to prevent dryness of the eyes. The lateral sides of the mice were then carefully shaved and aseptically prepared. Matched skin incisions were made to the shaved sides and the knee and elbow joints were tied together with nonabsorbable sutures (Reli no. SK7772) to facilitate coordinated movement. Surgical wound clips (BD Autoclips 9mm, no. 427631) and absorbable sutures (Ethicon no. J385H) were then used to join the skins together. Upon surgery, parabionts were injected subcutaneously with the anti-inflammatory Baytril (5 mg/kg) to prevent infection, and with the analgesics Carprofen (10mg/kg) and Buprenorphine (0.1mg/kg) to manage pain (single injection every 12 hours for up to 3 days post-surgery). In each injection, 0.5ml pre-warmed 0.9% (w/v) sodium chloride was also provided to prevent dehydration. Pairs were then kept in clean cages, and placed onto heating pads for up to 24 post-surgeries to maintain body temperature. Throughout the surgery and post-operative recovery each pair was monitored continuously and potential signs of pain and distress were recorded, while several physical characteristics were also analyzed, including pairs weights. After 15 days post-surgery, pairs were sedated again briefly to allow removal of the surgical wound clips and remnants of the absorbable sutures. Parabiotic pairs were maintained for 4-5 weeks (mean 31.5 days; Supplementary Fig 2) before processing for tissue collection and subsequent analysis.

### Blood chimerism analysis

To evaluate blood cross-circulation we followed the same approach as previously described^27^. Specifically, in the heterochronic pairs we used young mice carrying the congenic marker CD45.1 (JAX no. 002014) and old mice carrying the congenic marker CD45.2 (JAX no. 000664), while in the young isochronic pairs we used mice carrying either CD45.1 or CD45.2. For the blood chimerism analysis, spleens were extracted from the mice and single cells were mechanically isolated by passing the spleen through a 40uM filter. Erythrocytes were lysed with ACK lysis buffer (Thermo Fisher Scientific no. A1049201) for 3 minutes on ice and single cells were resuspended in HBSS (Thermo Fisher Scientific no. 14025-134) containing 2% FBS. Splenocytes were then filtered through a 40μM filter and stained with an antibody cocktail [Pacific Blue anti-CD45.1 (Biolegend no. 110722; 1:100 dilution), APC anti-CD45.2 (Biolegend no. 109814; 1:100 dilution), PE anti-TER-119 (Thermo Fisher Scientific, no. 12-5921-82; 1:200)], and the fixable viability dye Zombie Aqua (Biolegend no. 423101; 1:300 dilution) for 30 minutes on ice in the dark. Cells were then washed and fixed in 1% paraformaldehyde prior to analysis. Cells were gated on physical parameters to identify singlets followed by gating on the Zombie Aqua^low^TER-119^-^ population to identify live non-erythroid cells. These cells were subsequently gated as CD45.1^+^ or CD45.2 to measure the frequency of donor-derived blood cells from one partner in the spleen of the other partner. Flow cytometry analysis was performed using a BD LSR II flow cytometer (BD Biosciences) and data were analyzed with FlowJo software (version 10). We found that the partner-derived cells represented 30-50% of splenocytes (mean 41.3%; Supplementary Fig 1), consistent with successful establishment of parabiotic cross-circulation^45^. This analysis could not be applied to old isochronic mice, as old mice carrying the congenic marker CD45.1 were not available for purchase, however, the cross-circulation in old isochronic mice has been well-characterized previously^25^.

### Brain tissue dissociation

Brain tissue collection was performed at the same time of day (9-10am) processing one pair of mice per day, thus limiting circadian variation^135^. Brain tissue dissociation was performed as previously described^42^. Briefly, mice were CO_2_-anesthetized and then rapidly decapitated. Brains were extracted, and hindbrain regions were removed. The remaining tissue was mechanically and enzymatically dissociated into single cells and kept on ice for no longer than 1 hour until further processing.

### Single-cell RNA-sequencing

For the scRNA-seq experiments, 8 young (YX), 9 young heterochronic (YO), 9 young isochronic (YY), 8 old (OX), 11 old heterochronic (OY), and 12 old isochronic (OO) brains were analyzed, with 2 animals (1 pair) sacrificed per day as mentioned above. Briefly, after dissociation, cells were diluted in ice-cold PBS containing 0.4% BSA at a density of 1,000 cells/μl. For every sample, 17,400 cells were loaded into a Chromium Single Cell 3’ Chip (10x Genomics) and processed following the manufacturer’s instructions. Single-cell RNA-seq libraries were prepared using the Chromium v2 Single Cell 3’ Library & Gel Bead kit v2 and i7 Mutiplex kit (10X Genomics). Libraries were pooled based on their molar concentrations. Pooled libraries were then loaded at 2.07 pM and sequenced on a NextSeq 500 instrument (Illumina) with 26 bases for read1, 57 bases for read2 and 8 bases for Index1. Cell Ranger (version 1.2) (10X Genomics) was used to perform sample de-multiplexing, barcode processing and single cell gene unique molecular identifier (UMI) counting, while a digital expression matrix was obtained for each experiment with default parameters, mapped to the 10x reference for mm10, version 1.2.0. After the initial sequencing, the samples in each pool were re-pooled based on the actual number of cells detected by Cell Ranger (Supplementary Fig 2a), aiming to sequence each sample to a similar depth (number of reads/cell) (mean 43,107 reads/cell, Supplementary Fig 3c). Multiple NextSeq runs were conducted to achieve over 70% sequencing saturation as determined again by Cell Ranger (median: 75%).

### Raw data processing and quality control for cell inclusion

Basic processing and visualization of the scRNA-seq data were performed using the Seurat package (version 3.2.1.9002) ^136^ in R (version 3.6.1). The initial dataset contained 158,767 cells with data for 21,876 genes (Supplementary Fig 3). The average number of UMI (nCount_RNA) and non-zero genes (nFeature_RNA) are 2828.298 and 1206.153 respectively. The data were log normalized and scaled to 10,000 transcripts per cell. Variable genes were identified with FindVariableFeatures() function with the following parameters to set minimum and maximum average expressions and minimum dispersion: mean.cutoff(0.00125, 3), dispersion.cutoff(1, Inf). Next, principal component analysis (PCA) was carried out, and the top 50 PCs were stored. Clusters were identified with FindNeighbors() by constructing a K-nearest neighbor (KNN) graph, and clustered with the Louvain algorithm with FindClusters() at resolution 2, represented by UMAP projection. All clusters with only one cell were removed, and clusters with over 8% mitochondrial genes, clusters with min nCount_RNA less than 1000, and clusters with min nFeature_RNA 500 were flagged for exclusion, resulting in 80 initial clusters. Animals with low average number of genes above 0 (<700), percentage mitochondria > 1.5, and not having cell contribution to each cluster were compared to surgery notes and assessed for exclusion. In total, 5 isochronic old and 1 isochronic young were removed from the dataset, and the above steps were clustering steps were performed at resolution 2. For quality control (QC) filtering, we selectively removed clusters with minimum percent mitochondria 0, maximum percent mitochondria 5%, min_nFeature_RNA 250, max_nFeature_RNA 6000, min_nCount_RNA 200, max_nCount_RNA 30000, min_cells=5. After the second round QC, we retained 130,889 cells and 20,905 genes. The average nCount_RNA, non-zero genes, percent mitochondrial RNA, and percent ribosomal RNA were 2736.187, 1368.007, 1.149, and 5.135. We re-clustered at resolution 2, to identify 69 clusters. The final pre-processing step was to remove likely doublet artifacts arising from the co-capture of multiple cells in one droplet. After an initial round of cluster identity determination as assessed in the next section, we employed a doublet finding technique by searching for the top differential markers of each identified cluster/sub-cluster with the FindMarkers() function, and marked doublets/multiplets as any cluster in which >40% of its cells express 7 of the top 10 genes specific to an initially identified cell type and any other outside of the class of the cell type it is associated with. These clusters were removed from the downstream analysis, and clustering was again performed at resolution 2, representing 105,329 cells and 69 clusters across 20,905 genes.

### Determination of cell type identity

We used multiple cell-specific/enriched gene markers that have been previously described in the literature to assist in determining cell type identity^42^. We then arranged all the identified cell types based on their expression profile, lineage, function, and topology into 5 classes of cells. For each group, we re-clustered the sub-categorized cell types using top 50 PCs at resolution 5. The annotation of the sub-clusters was performed similar to the identification of the main cell clusters.

### Differential gene expression analysis

After initial quality control pre-processing and determination of cellular identities, we utilized the muscat package (version 1.0.0) in R (3.6.1) to perform pseudobulk differential gene expression (DGE) analysis with edgeR^54–56^. Seurat objects were exported to SingleCellExperiments, and reads were collapsed per animal to “sum” based on “counts”. The “rejuvenation framework” RJV follows the design contrast (OY-OX)-(OO-YX), assigning in the design matrix: OY: 1, OO:-1, OX:-1, YX: 1. The “aging acceleration framework” AGA follows the design contrast (YO-YX)-(YY-OX), assigning in the design matrix: YO: 1, YY:-1, YX:-1, OX: 1. Pairwise comparisons for OXvYX, OYvOX, OYvOO, OOvOX, YOvYX, YOvYY, YYvYX were also computed. Via muscat, edgeR generates a logFC, logCPM, F, p_val, p_adj.loc, p_adj.glb. We used the Benjamini-Hochberg adjusted q-value p_val.loc in all downstream thresholding. Our ability to establish a baseline level of transcription is reliant on the number of cells measured and thus larger clusters’ variation can be more adequately modelled. HypEPC and TNC did not contain enough cells over multiple animals to successfully derive statistics. For all mouse types, raw normalized TPM values were calculated, and percentage of expression per animal type. For heatmap representations, the log2 z-score for the rejuvenation or aging-acceleration process, mice were calculated gene-wise (by row). For further comparison, we completed MAST analysis at the single-cell resolution level as part of the Seurat framework FindMarkers()^136,137^. Per cluster, pairwise comparisons for OXvYX, OYvOX, OYvOO, OOvOX, YOvYX, YOvYY, YYvYX were computed. To retain DGE information for the maximal amount of genes, we used the parameters: min.cells.group = 1, min.cells.feature = 1, min.pct = 0, logfc.threshold = 0, only.pos = FALSE. Benjamini-Hochberg FDR was computed.

### Pathway analysis

Gene Set Enrichment Analysis (GSEA) was performed with the fgsea R package (version 1.12.0) ^138^. Using the protocol previously implemented^139^, for each cell population and DGE comparison, genes were ranked by multiplying −log10(p_val) with the sign of the logFC and converted to *Homo sapiens* orthologs using biomaRt (version 2.46.2). From MSigDB, we used 5 gene sets: *Hallmark pathways, GO Biological Process, KEGG, BioCarta*, and *Reactome*. In fgsea, 1000 permutations were performed with minimum gene set size 15, max 500. Gene sets with FDR≤0.25 were considered significantly enriched. Term annotations and grouping of those overrepresenting the same pathway were derived from Cytoscape software (version 3.5.1) and the AutoAnnotate app (version1.2) as previously described^42^. The Normalized Enrichment Scores (NES) directionality was used to collate cell type pathways per DGE comparison. Dot plot representations are a composite of FDR and NES.

### Gene regulatory networks

We employed SCENIC to assess gene regulatory networks (GRNs) and score their activity, using the R implementation (version 1.1.2-2) ^70^. Each animal type was analyzed with respect to each lineage. Briefly, we used GENIE3 to identify genes that are co-expressed with transcription factors. Then, RCisTarget prunes these co-expression modules to create GRNs (regulons). The direct targets of each transcription factor are found using cis-regulatory motif analysis. AUCell scores each regulon’s activity, binarized to on/off at threshold 0.7. The regulon activity per animal type per lineage is graphed with rose diagram histograms. Regulons whose TFs change logFC direction between RJV and AGA are identified, and those regulons changing direction in at least 8 cell clusters are further investigated. The magnitude of the logFC difference between RJV and AGA can take the form of the absolute difference, with the sign of the difference as positive if rejuvenation is > 0 and aging acceleration is <0, negative if aging acceleration is > 0 and rejuvenation is < 0, or zeroed out if they are in the same direction. Regulon matrices of each animal type across all clusters, along with row-wise regulon count, are reported.

### Cell-cell communication

Cell-cell communication between cell types per animal type was assessed using the CellChat tool^81^. Number of interactions graphs per animal were thresholded at interactions reaching p ≤ 0.05 and graphed with netVisual_circle. Rejuvenation-associated construct graphs were the subset of unique receptor:ligand:source:target combinations of interactions occurring only in OY and YX, and aging-associated construct graphs were the same combination occurring only in YO and OX, graphed via the circlize package (0.4.13.1001)^140^. EC receptors in all 6 animal types were collapsed into a master list, with DGE padj/logFC graphed via ggplot2. EC receptor graphs per animal type were constructed via CellChat netVisual_aggregate.

Cell-cell communication per comparison was also conducted as previously described^42^ using the CCInx package (0.5.1). Per comparison plots were generated between ligand:receptor pairs using the CCInx tool.

### Cellular senescence

Cellular senescence was investigated using functional enrichment on pre-ranked genes against known senescence marker genes as described in the literature ^107,108^. Briefly, the pre-ranked (-log10(pval) multiplied by the sign of the logFC) Aging, RJV, and AGA gene lists were permuted 1000 times against the gene set using the fgsea algorithm implemented in ClusterProfiler (version 3.14.3). The Normalized Enrichment Score (NES) and adjusted p-value (Benjamini-Hochberg) are reported.

### Endothelial cell class assignment

Endothelial cells “zonation” was assessed through deep learning using the CellAssign framework^141^ (version 0.99.21, tensorflow_2.2.0.9000). Gene markers from Zhao et al. 2020^50^ and other sources^48–50^ were used to define arterial-venous-capillary markers. The learning rate used was 1e-2, with a min_delta of 0.25, and 10 runs on a V100 GPU hosted on the FAS Cannon cluster.

### RNA *in situ* hybridization

RNA *in situ* hybridization was performed on fresh-frozen brain tissue from at least three mice of each relevant condition (YX, OY, OO, OX). For sample preparation, mice were sacrificed via cervical dislocation, and brains were rapidly extracted and embedded in OCT (Tissue Tek) on dry ice, and subsequently stored at −80 °C until further processing. Brains were divided into 14-μm cryostat sections and RNA *in situ* hybridizations were carried out using the RNA *in situ* Multiplex Fluorescent Manual Assay kit (Advanced Cell Diagnostics, ACD) per manufacturer’s instructions. Briefly, thawed sections were fixed in 4% paraformaldehyde in PBS and dehydrated in sequential incubations with ethanol, followed by 30-minute Protease IV treatment and washing in 1X PBS. Appropriate combinations of hybridization probes were incubated on tissue for 2 hours at 40°C, followed by four amplification steps. Sections were subsequently stained with DAPI and mounted with Prolong Gold mounting medium (Thermo Fisher Scientific no. P36930). Brain regions were selected based upon areas of high expression levels of assessed examined genes, according to the Allen Brain Atlas^142^. Commercially available and validated probes for *Cdkn1a* (ACD no. 408551), *Hspa1a* (ACD no. 488351), Klf6 (ACD no. 426901), and *Pecam1* (ACD no. 316721) were utilized per manufacturer’s instructions. For each mouse and tissue, 3 bregma-matched sections were imaged. Images (4 per tissue section) were acquired with a Zeiss LSM 880 Confocal Microscope with identical settings across sections and represented as maximum intensity projections of acquired confocal z-stacks. Analysis was done utilizing a script within CellProfiler software (v.3)99 in which *Cdkn1a, Hspa1a*, or *Klf6* puncta with diameter between 1 and 12 pixels located within the perinuclear space (100 pixels of DAPI-positive nuclei) were identified and quantified. Cells with ≥2 *Pecam1+* puncta were designated *Pecam1+* endothelial cells (ECs). For *Hspa1a* and *Cdkn1a* experiments, EC marker *Pecam1* was labeled by fluorophore Atto 647, while target probes were labeled by Atto 488 (Hspa1a) and Atto 550 (*Cdkn1a*). For *Klf6* experiments, *Pecam1* was labeled by fluorophore Alexa 488 while *Klf6* was labeled by fluorophore Atto 550. Lipofuscin granules largely associated with aged brain tissue were avoided utilizing the 1-12-pixel cutoff for identifying puncta. For each animal, an unstained tissue was imaged as a negative control and to assess levels of background fluorescence.

### Statistical tests

No statistical methods were used to predetermine sample sizes; our samples sizes were determined iteratively. No randomization was performed. Data collection and analysis were not performed blind to the conditions of the experiments. All statistical analyses were performed with R (v3.6.0) or GraphPad Prism (v.7.04). To generate p-values for cell counts, ANOVA was conducted between animal types per cell type (rstatix 0.6.0). For validation of gene expression changes by RNA *in situ* hybridization, Mann-Whitney U test or Student’s t-test were conducted as indicated (rstatix 0.6.0).

