## Supplementary Files for "Heterochronic parabiosis reprograms the mouse brain transcriptome by shifting aging signatures in multiple cell types"

Fig_S1

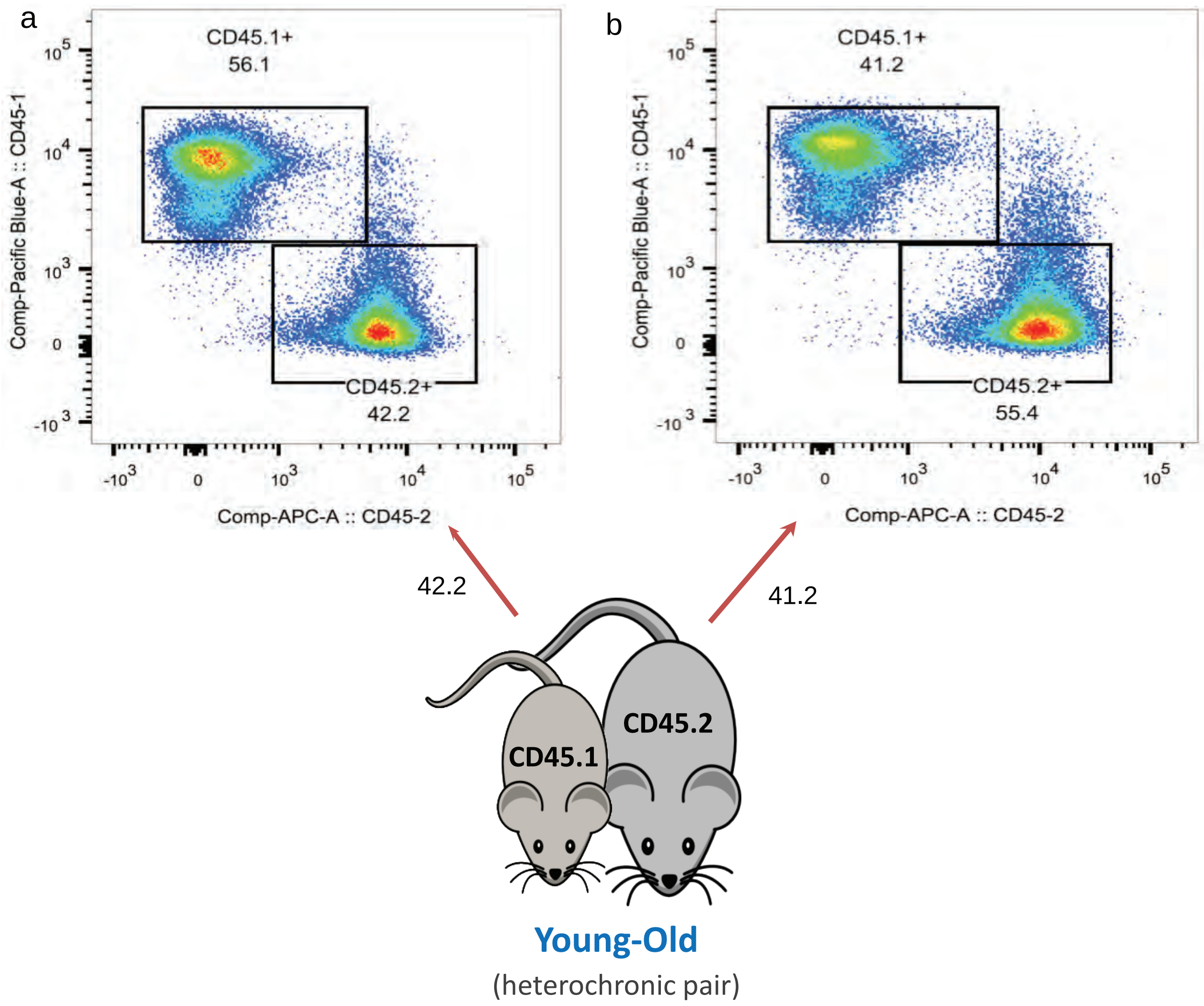
a
CD45.1+
56.1
CD45.2+
42.2
Comp-Pacific Blue-A :: CD45-1
Comp-APC-A :: CD45-2
b
CD45.1+
41.2
CD45.2+
55.4
Comp-Pacific Blue-A :: CD45-1
Comp-APC-A :: CD45-2
42.2
41.2
CD45.1
CD45.2
Young-Old
(heterochronic pair)

#### Supplementary Figures

**Supplementary Fig 1**. **Confirmation of blood chimerism.** Representative flow cytometry analysis of CD45.1 and CD45.2 expression markers on splenocytes isolated from (**a**) young (CD45.1$^{+}$) and (**b**) old mice (CD45.2$^{+}$) following heterochronic parabiosis. In each plot, the percentage of donor-derived blood cells from one partner in the spleen of the other partner is depicted by arrows.

Fig_S2

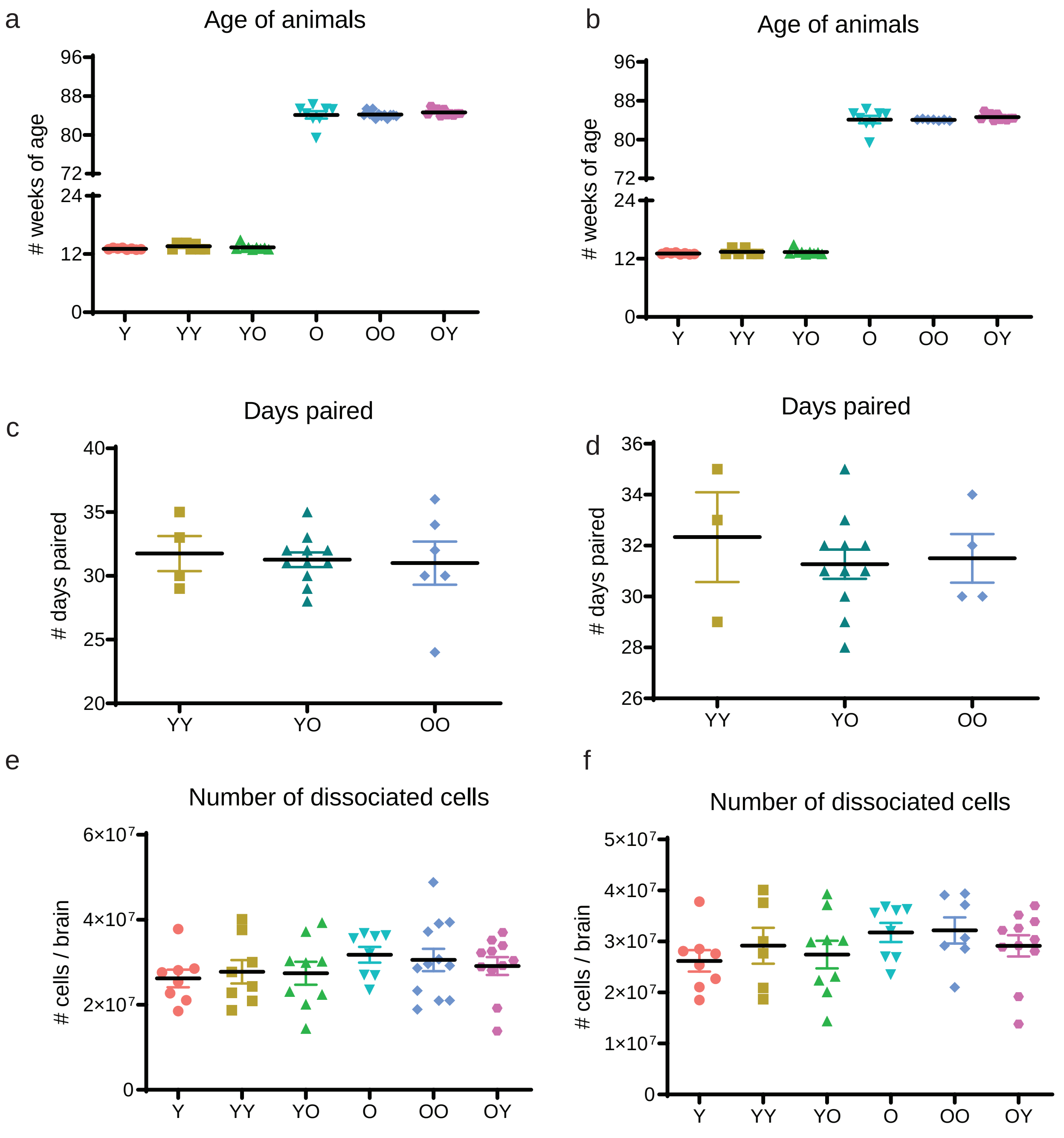
a
Age of animals
### weeks of age
96
88
80
72
24
12
0
Y
YY
YO
O
OO
OY
b
Age of animals
### weeks of age
c
Days paired
### days paired
40
35
30
25
20
d
Days paired
### days paired
36
34
32
30
28
26
e
Number of dissociated cells
### cells / brain
6×10^7
4×10^7
2×10^7
f
Number of dissociated cells
### cells / brain
5×10^7
4×10^7
3×10^7
2×10^7
1×10^7

**Supplementary Fig 2. Sample metrics.** Profiling of animals and their derived brain cells used for sequencing, before (a,c,e) and after (b,d,f) quality control filtering in which certain animals were omitted (see Methods). **a-b.** Age of mice in weeks prior to parabiosis surgeries. **c-d.** Number of days joined across parabiotic pairs. **e-f.** Number of dissociated cells analyzed per brain across all animal types.

Fig_S3

a

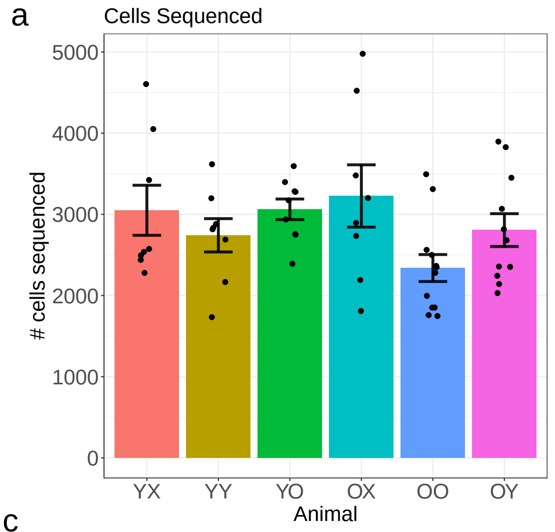


b

| Animal | # Mice | # Cells |
|---|---|---|
| YX | 8 | 24401 |
| YY | 8 | 21935 |
| YO | 9 | 27557 |
| OX | 8 | 25811 |
| OO | 12 | 28066 |
| OY | 11 | 30865 |

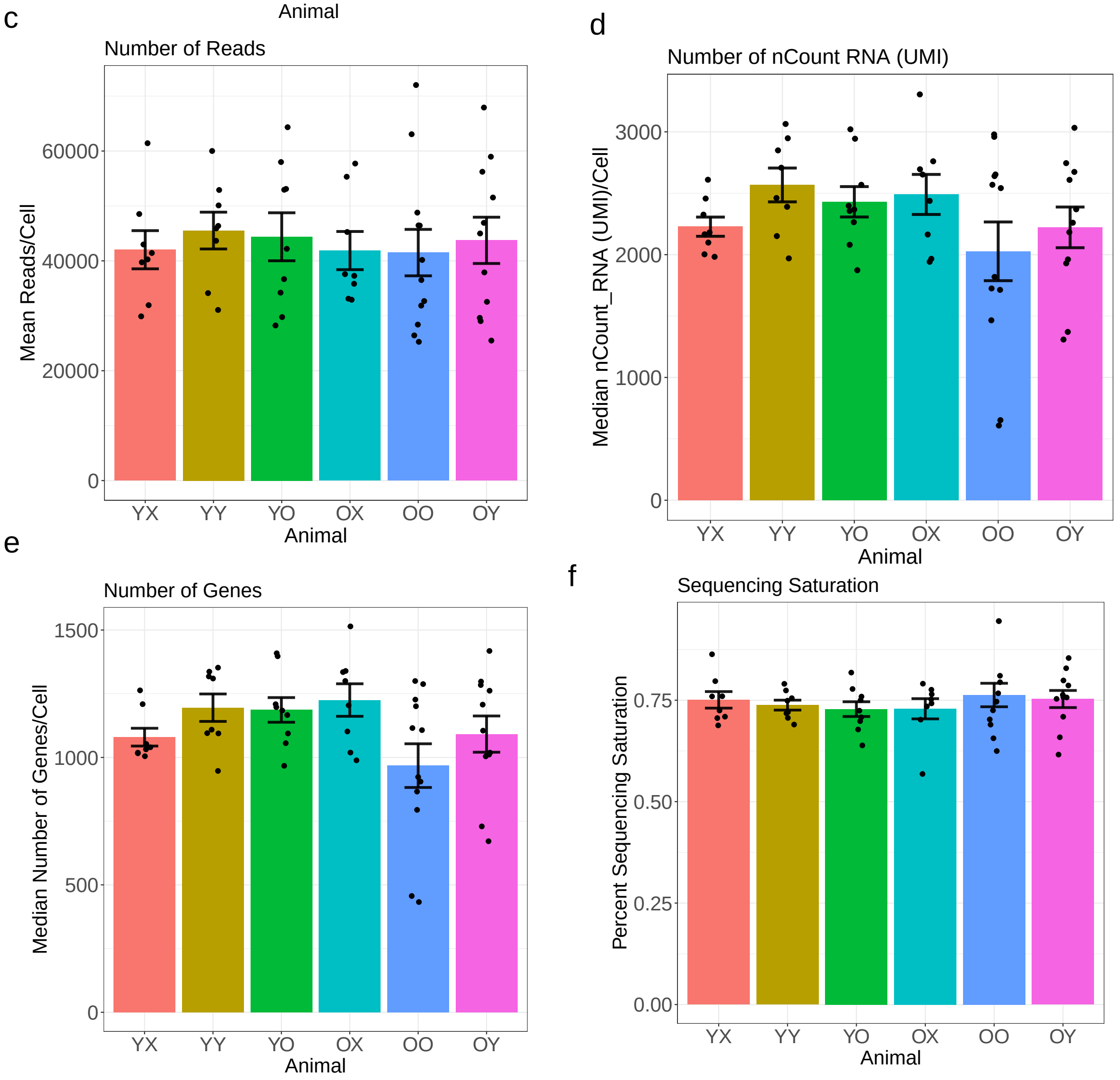

**Supplementary Fig 3. Sequencing metrics.** Scatter plots showing metrics of all sequenced young and old cells prior to any cell filtering: **a.** Number of cells sequenced by animal. **b.** Total number of animals and cells analyzed. **c.** Mean number of mapped reads per cell by animal. **d.** Median number of nCount RNA (UMI) detected per cell by animal. **e.** Median number of genes detected per cell by animal. Data presents mean or median ± SEM.

Fig_S4

a

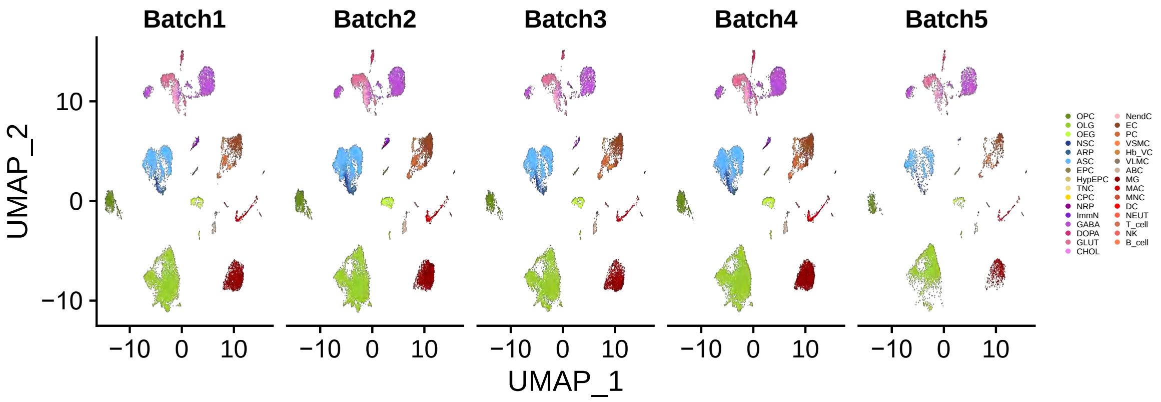
Batch1
Batch2
Batch3
Batch4
Batch5
UMAP_2
10
0
−10
−10 0 10
UMAP_1
OPC
OLG
OEG
NSC
ARP
ASC
EPC
HypEPC
TNC
CPC
NRP
ImmN
GABA
DOPA
GLUT
CHOL
NendC
EC
PC
VSMC
Hb_VC
VLMC
ABC
MG
MAC
MNC
DC
NEUT
T_cell
NK
B_cell

b

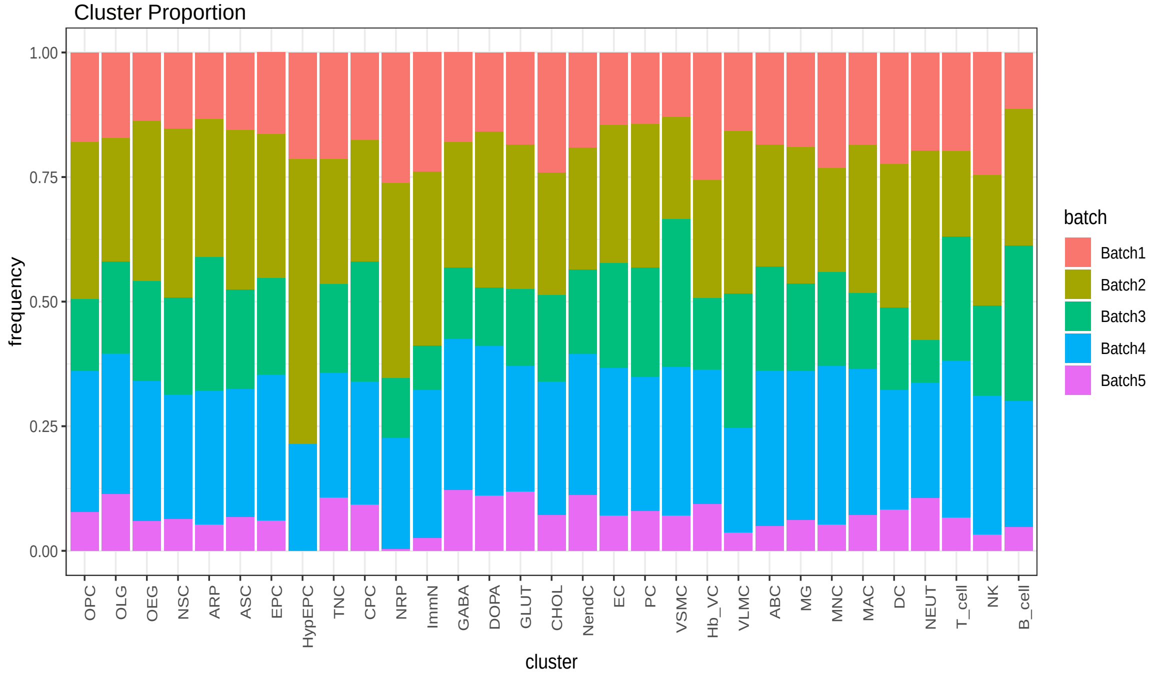
Cluster Proportion
frequency
1.00
0.75
0.50
0.25
0.00
OPC
OLG
OEG
NSC
ARP
ASC
EPC
HypEPC
TNC
CPC
NRP
ImmN
GABA
DOPA
GLUT
CHOL
NendC
EC
PC
VSMC
Hb_VC
VLMC
ABC
MG
MNC
MAC
DC
NEUT
T_cell
NK
B_cell
cluster
batch
Batch1
Batch2
Batch3
Batch4
Batch5

**Supplementary Fig 4. Distribution of 50 animals across 5 sequencing batches, with respect to cell clusters**. **a**. UMAP projection of color-coded batches over clusters that passed filtering criteria. **b.** Frequency of each color-coded batch representation in each cell type. All cell types are represented by cells from all batches, except for HypEPC in batch 5, probably due to its small size.

Fig_S5

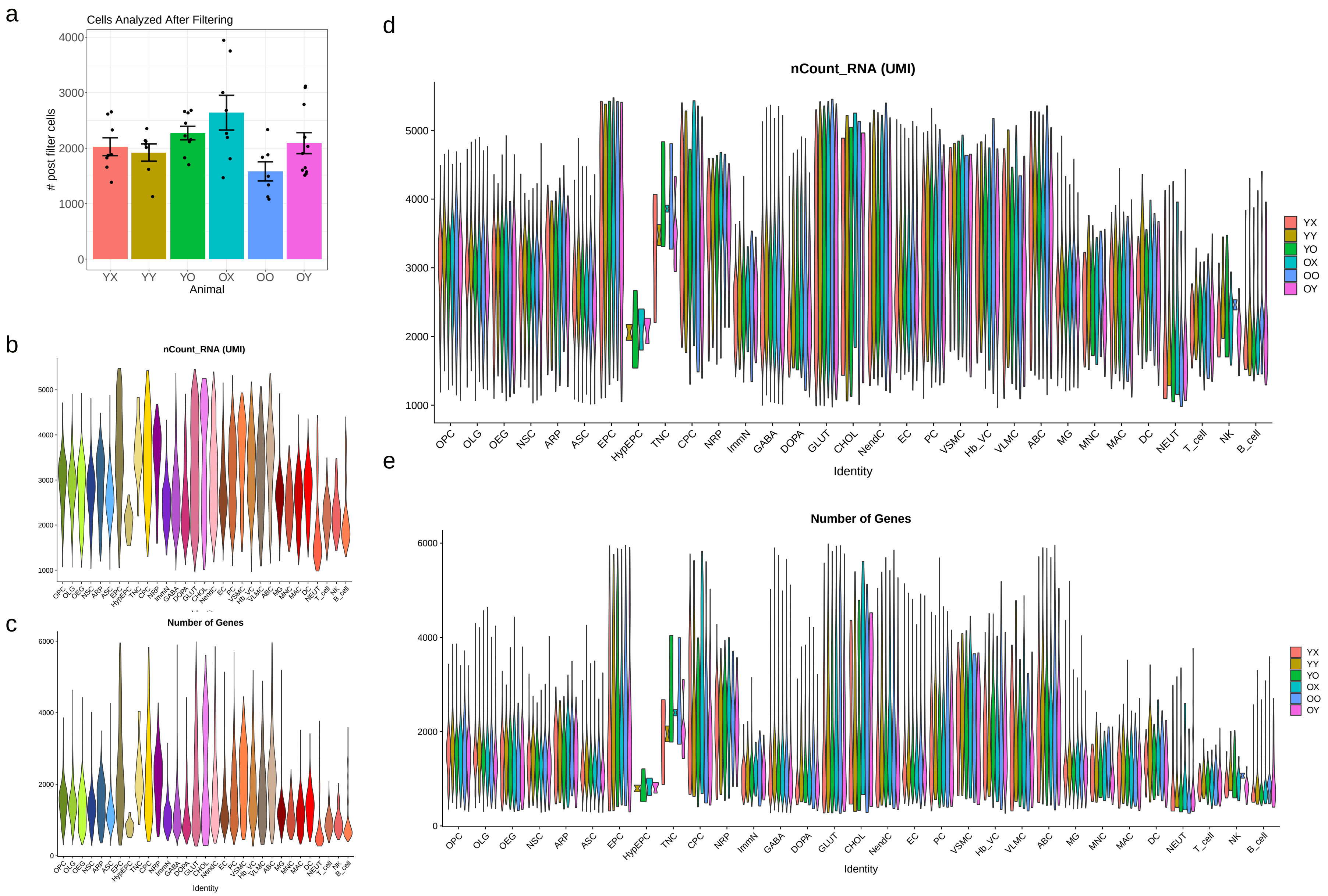
a
Cells Analyzed After Filtering
### post filter cells
Animal
YX
YY
YO
OX
OO
OY
b
nCount_RNA (UMI)
c
Number of Genes
Identity
d
nCount_RNA (UMI)
Identity
e
Number of Genes
Identity
OPC
OLG
OEG
NSC
ARP
ASC
EPC
HypEPC
TNC
CPC
NRP
ImmN
GABA
DOPA
GLUT
CHOL
NendC
EC
PC
VSMC
Hb_VC
VLMC
ABC
MG
MNC
MAC
DC
NEUT
T_cell
NK
B_cell

**Supplementary Fig 5. Primary data analysis. a**. Bar plot showing the number of cells analyzed by animal after cell filtering, in which all cells were successfully assigned to a specific cell type (data presents mean ± SEM). **b-e.** Violin plots showing QC metrics, plots in (b, c) showing aggregated data of cells of all brain types, while plots in (d, e) showing individual cell data separated by animal type: (b, d) showing nCount RNA (UMI) per cell type. (c, e) showing nFeature RNA (number of unique genes) detected per cell.

Fig_S6

a

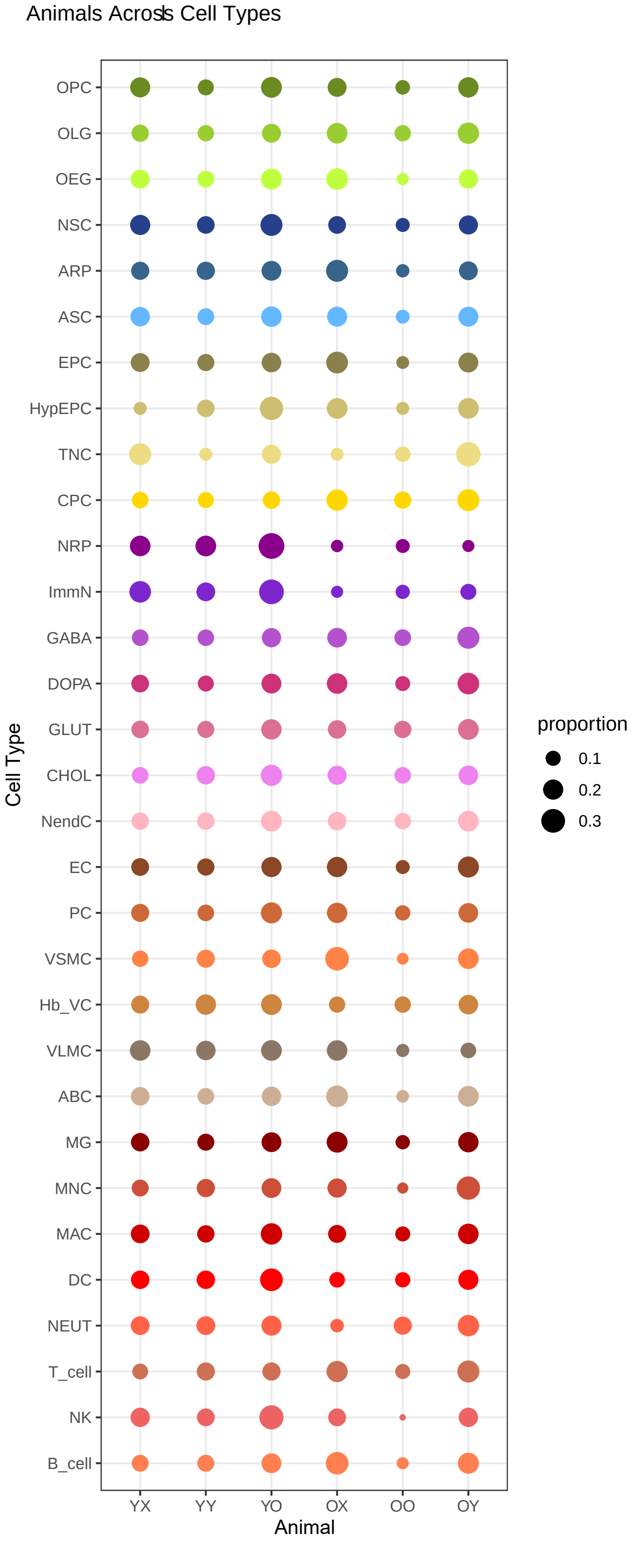
Animals Across Cell Types
OPC
OLG
OEG
NSC
ARP
ASC
EPC
HypEPC
TNC
CPC
NRP
ImmN
GABA
DOPA
GLUT
CHOL
NendC
EC
PC
VSMC
Hb_VC
VLMC
ABC
MG
MNC
MAC
DC
NEUT
T_cell
NK
B_cell
Cell Type
YX
YY
YO
OX
OO
OY
Animal
proportion
0.1
0.2
0.3

b

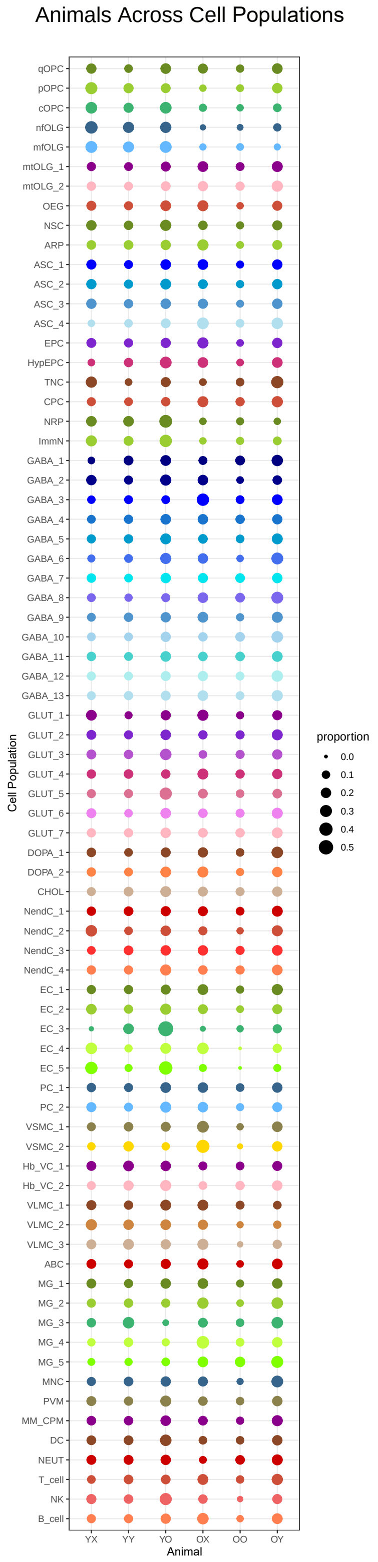
Animals Across Cell Populations
qOPC
pOPC
cOPC
nfOLG
mfOLG
mtOLG_1
mtOLG_2
OEG
NSC
ARP
ASC_1
ASC_2
ASC_3
ASC_4
EPC
HypEPC
TNC
CPC
NRP
ImmN
GABA_1
GABA_2
GABA_3
GABA_4
GABA_5
GABA_6
GABA_7
GABA_8
GABA_9
GABA_10
GABA_11
GABA_12
GABA_13
GLUT_1
GLUT_2
GLUT_3
GLUT_4
GLUT_5
GLUT_6
GLUT_7
DOPA_1
DOPA_2
CHOL
NendC_1
NendC_2
NendC_3
NendC_4
EC_1
EC_2
EC_3
EC_4
EC_5
PC_1
PC_2
VSMC_1
VSMC_2
Hb_VC_1
Hb_VC_2
VLMC_1
VLMC_2
VLMC_3
ABC
MG_1
MG_2
MG_3
MG_4
MG_5
MNC
PVM
MM_CPM
DC
NEUT
T_cell
NK
B_cell
Cell Population
YX
YY
YO
OX
OO
OY
Animal
proportion
0.0
0.1
0.2
0.3
0.4
0.5

**Supplementary Fig 6. Representation of each animal type's distribution within each cell type**. **a.** Dot plot representation of each cell type's representation by each animal type. Size of the dot is proportional to the number of cells contributed by each animal type within a cell type. **b**. Dot plot representation of each subpopulation's representation by each animal type. Size of the dot is proportional to the number of cells contributed by each animal type within a subpopulation.

Fig_S7

a

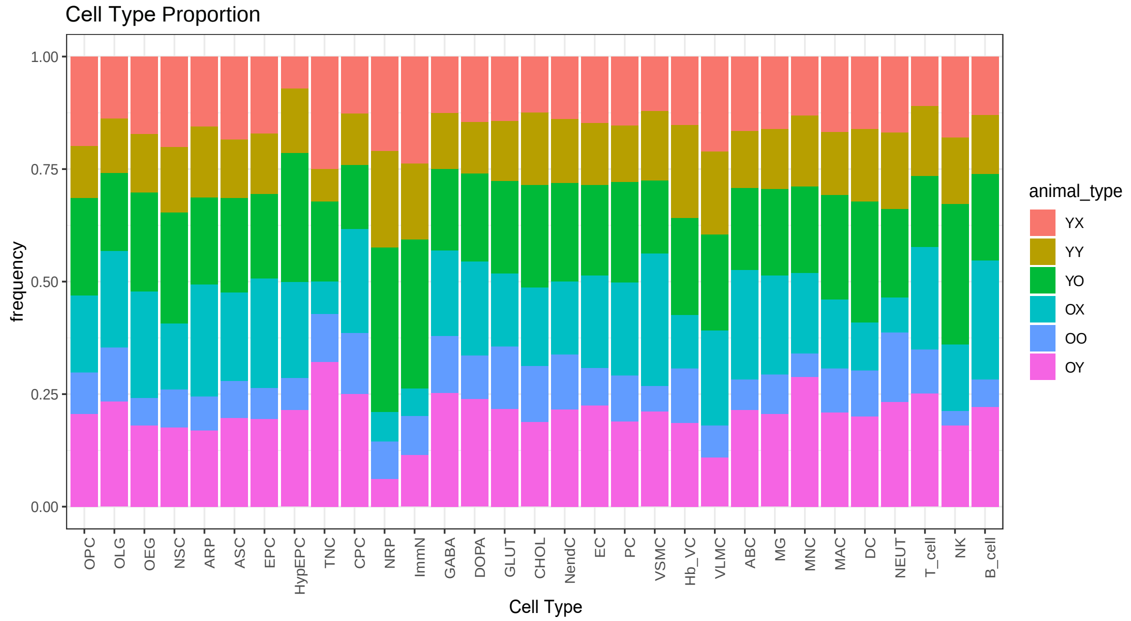
Cell Type Proportion
frequency
1.00
0.75
0.50
0.25
0.00
OPC
OLG
OEG
NSC
ARP
ASC
EPC
HypEPC
TNC
CPC
NRP
ImmN
GABA
DOPA
GLUT
CHOL
NendC
EC
PC
VSMC
Hb_VC
VLMC
ABC
MG
MNC
MAC
DC
NEUT
T_cell
NK
B_cell
Cell Type
animal_type
YX
YY
YO
OX
OO
OY

b

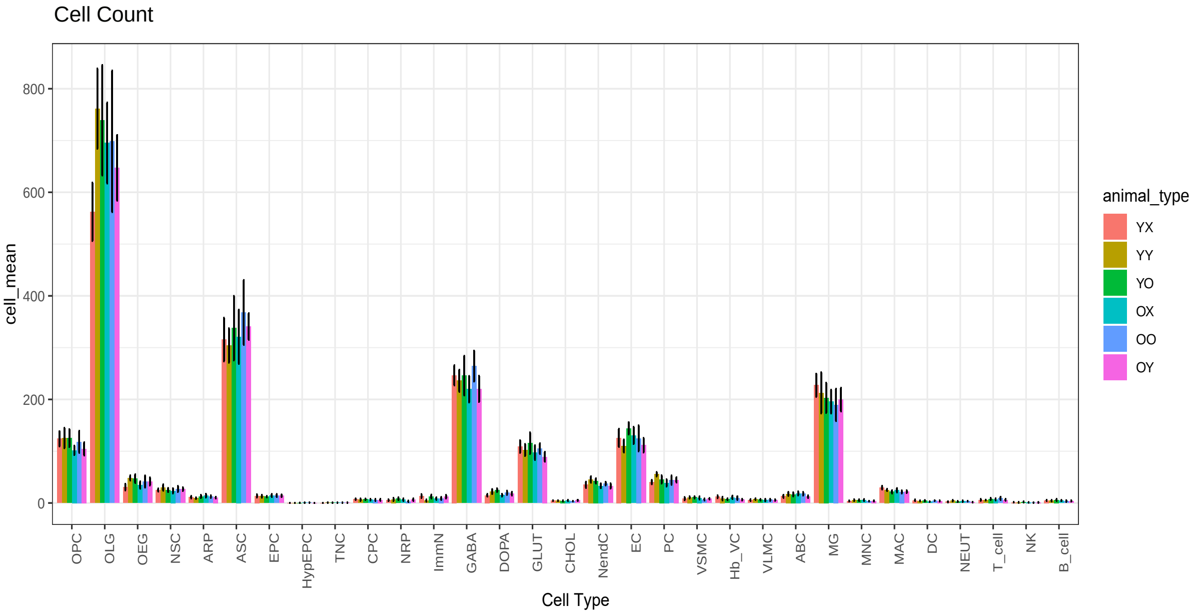
Cell Count
cell_mean
800
600
400
200
0
OPC
OLG
OEG
NSC
ARP
ASC
EPC
HypEPC
TNC
CPC
NRP
ImmN
GABA
DOPA
GLUT
CHOL
NendC
EC
PC
VSMC
Hb_VC
VLMC
ABC
MG
MNC
MAC
DC
NEUT
T_cell
NK
B_cell
Cell Type
animal_type
YX
YY
YO
OX
OO
OY

**Supplementary Fig 7. Cell type composition and cell count from each animal type. a.** Frequency bar plot demonstrating composition of each cell type with respect to animal type. **b.** Bar plot of raw cell counts, +/- SEM, with respect to each animal. All animals contribute to all cell types. ANOVA p-values for pairwise iterations can be found in Supplementary Table 2.

Fig_S8

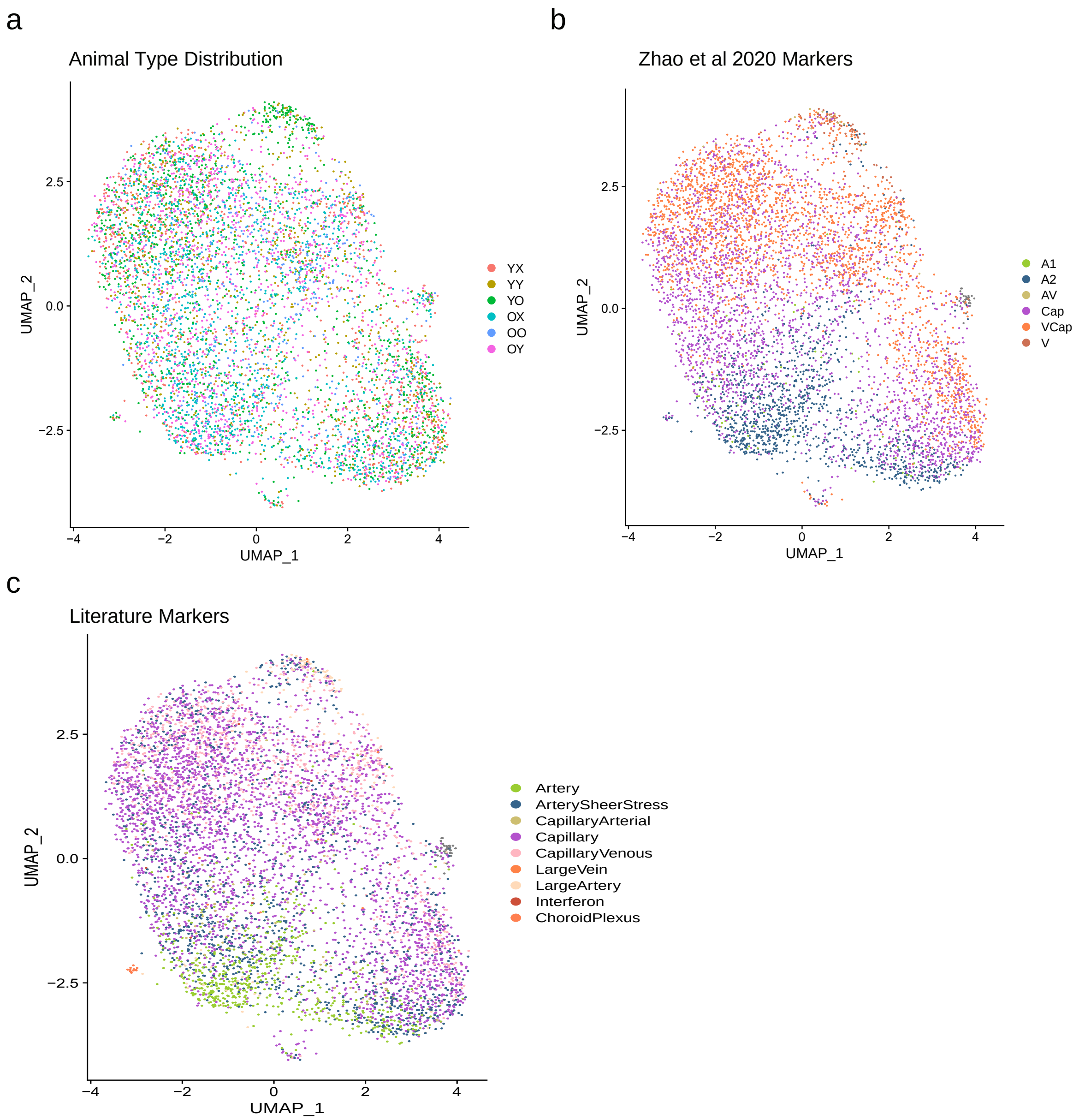
a
Animal Type Distribution
UMAP_2
UMAP_1
2.5
0.0
−2.5
−4
−2
0
2
4
YX
YY
YO
OX
OO
OY
b
Zhao et al 2020 Markers
A1
A2
AV
Cap
VCap
V
c
Literature Markers
Artery
ArterySheerStress
CapillaryArterial
Capillary
CapillaryVenous
LargeVein
LargeArtery
Interferon
ChoroidPlexus

**Supplementary Fig 8. Animal type distribution and machine learning approaches to explore EC arteriovenous zonation. a**. Animal type cell distribution across EC subclusters. **b-c.** Probabilistic programming cell class assignment using EC marker genes described by Zhao et al 2020[4] (b) and others[4,3,2] (c).

Fig_S9

a

Aging-RJV

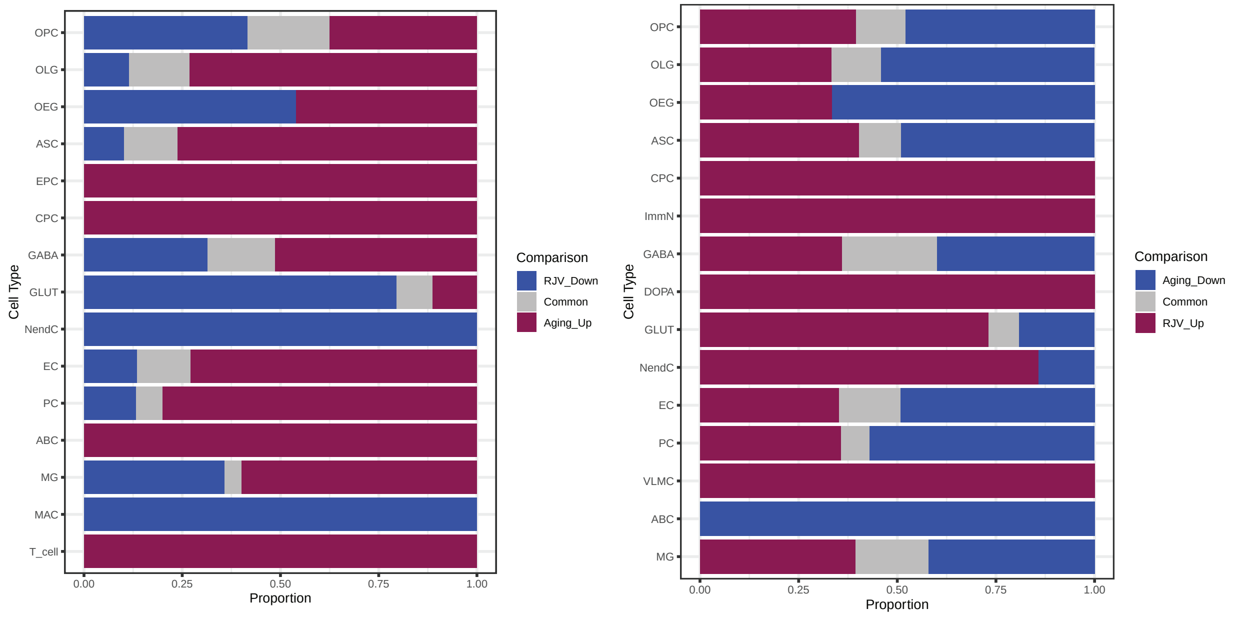


b

Aging-AGA

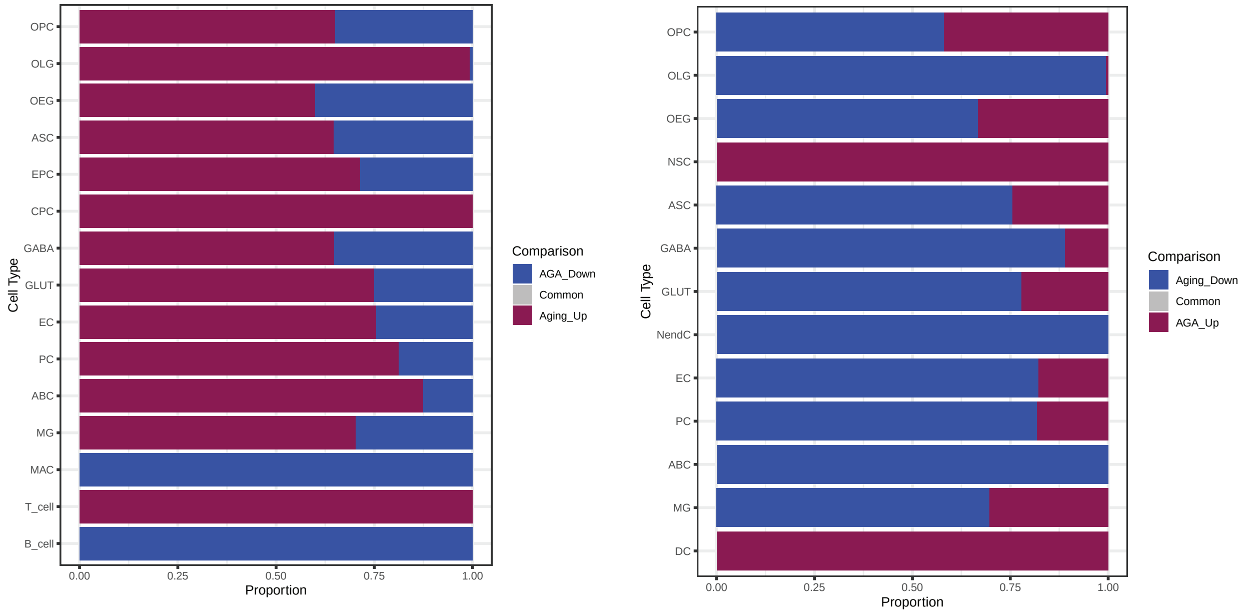

**Supplementary Fig 9. Composition of DGEs per cell type between Aging-RJV, and Aging-AGA. a.** Bar graph of each cell type's total FDR ≤ 0.05 DGEs split by logFC direction. The proportion of DGEs reflecting Aging and RJV is depicted, as well as the fraction of overlapping signatures (intersection in grey). **b.** Bar graph of each cell type's total FDR ≤ 0.05 DGEs split by logFC direction. The proportion of DGEs reflecting Aging and AGA is depicted, as well as the fraction of overlapping signatures (intersection in grey).

Fig_S10

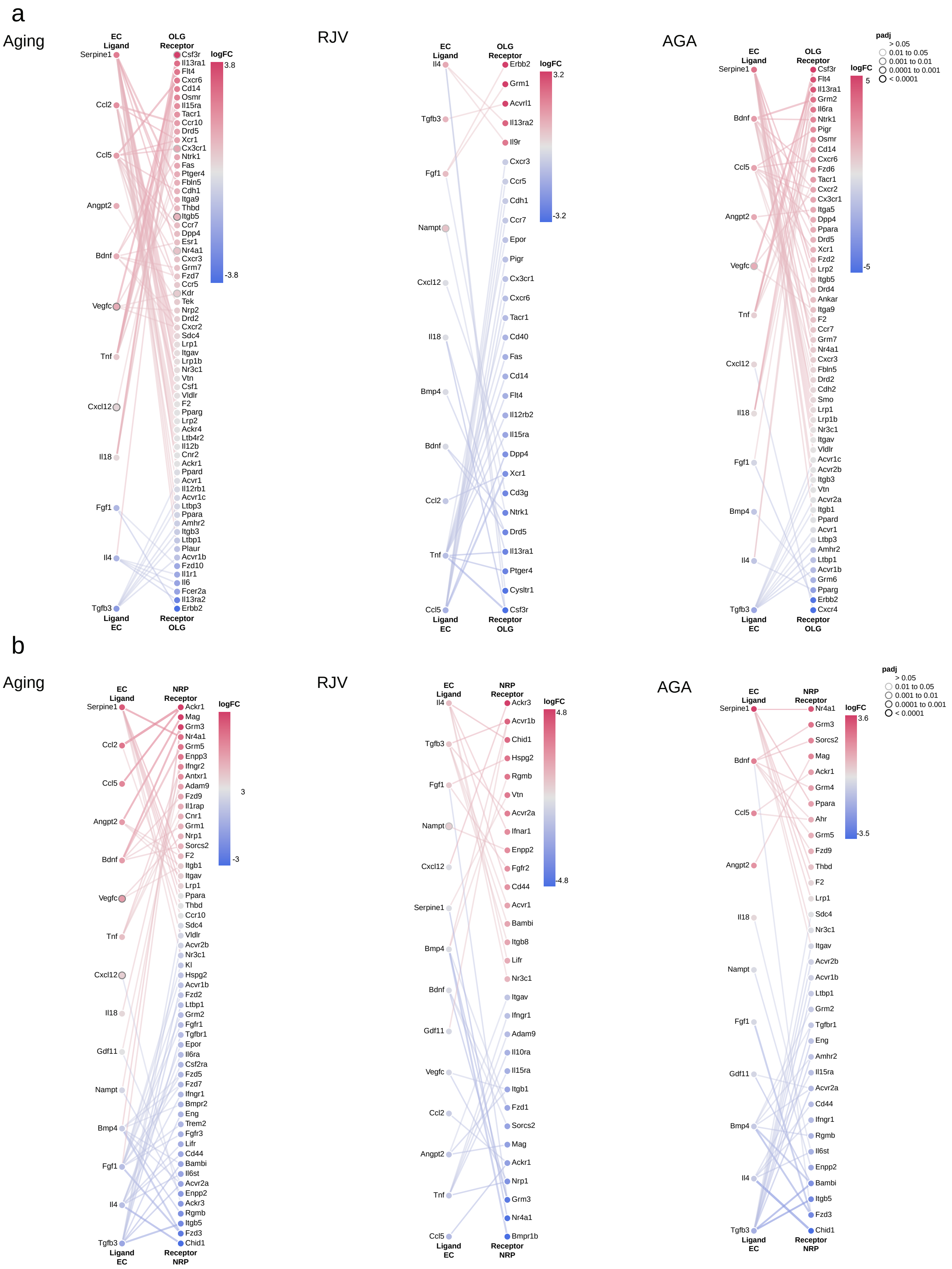
a
Aging
EC Ligand
OLG Receptor
logFC
3.8
-3.8
Serpine1
Ccl2
Ccl5
Angpt2
Bdnf
Vegfc
Tnf
Cxcl12
Il18
Fgf1
Il4
Tgfb3
Csf3r
Il13ra1
Flt4
Cxcr6
Cd14
Osmr
Il15ra
Tacr1
Ccr10
Drd5
Xcr1
Cx3cr1
Ntrk1
Fas
Ptger4
Fbln5
Cdh1
Itga9
Thbd
Itgb5
Ccr7
Dpp4
Esr1
Nr4a1
Cxcr3
Grm7
Fzd7
Ccr5
Kdr
Tek
Nrp2
Drd2
Cxcr2
Sdc4
Lrp1
Itgav
Lrp1b
Nr3c1
Vtn
Csf1
Vldlr
F2
Pparg
Lrp2
Ackr4
Ltb4r2
Il12b
Cnr2
Ackr1
Ppard
Acvr1
Il12rb1
Acvr1c
Ltbp3
Ppara
Amhr2
Itgb3
Ltbp1
Plaur
Acvr1b
Fzd10
Il1r1
Il6
Fcer2a
Il13ra2
Erbb2
Ligand EC
Receptor OLG
RJV
EC Ligand
OLG Receptor
logFC
3.2
-3.2
Il4
Tgfb3
Fgf1
Nampt
Cxcl12
Il18
Bmp4
Bdnf
Ccl2
Tnf
Ccl5
Erbb2
Grm1
Acvrl1
Il13ra2
Il9r
Cxcr3
Ccr5
Cdh1
Ccr7
Epor
Pigr
Cx3cr1
Cxcr6
Tacr1
Cd40
Fas
Cd14
Flt4
Il12rb2
Il15ra
Dpp4
Xcr1
Cd3g
Ntrk1
Drd5
Il13ra1
Ptger4
Cysltr1
Csf3r
Ligand EC
Receptor OLG
AGA
EC Ligand
OLG Receptor
padj
> 0.05
0.01 to 0.05
0.001 to 0.01
0.0001 to 0.001
< 0.0001
logFC
5
-5
Serpine1
Bdnf
Ccl5
Angpt2
Vegfc
Tnf
Cxcl12
Il18
Fgf1
Bmp4
Il4
Tgfb3
Csf3r
Flt4
Il13ra1
Grm2
Il6ra
Ntrk1
Pigr
Osmr
Cd14
Cxcr6
Fzd6
Tacr1
Cxcr2
Cx3cr1
Itga5
Dpp4
Ppara
Drd5
Xcr1
Fzd2
Lrp2
Itgb5
Drd4
Ankar
Itga9
F2
Ccr7
Grm7
Nr4a1
Cxcr3
Fbln5
Drd2
Cdh2
Smo
Lrp1
Lrp1b
Nr3c1
Itgav
Vldlr
Acvr1c
Acvr2b
Itgb3
Vtn
Acvr2a
Itgb1
Ppard
Acvr1
Ltbp3
Amhr2
Ltbp1
Acvr1b
Grm6
Pparg
Erbb2
Cxcr4
Ligand EC
Receptor OLG
b
Aging
EC Ligand
NRP Receptor
logFC
3
-3
Serpine1
Ccl2
Ccl5
Angpt2
Bdnf
Vegfc
Tnf
Cxcl12
Il18
Gdf11
Nampt
Bmp4
Fgf1
Il4
Tgfb3
Ackr1
Mag
Grm3
Nr4a1
Grm5
Enpp3
Ifngr2
Antxr1
Adam9
Fzd9
Il1rap
Cnr1
Grm1
Nrp1
Sorcs2
F2
Itgb1
Itgav
Lrp1
Ppara
Thbd
Ccr10
Sdc4
Vldlr
Acvr2b
Nr3c1
Kl
Hspg2
Acvr1b
Fzd2
Ltbp1
Grm2
Fgfr1
Tgfbr1
Epor
Il6ra
Csf2ra
Fzd5
Fzd7
Ifngr1
Bmpr2
Eng
Trem2
Fgfr3
Lifr
Cd44
Bambi
Il6st
Acvr2a
Enpp2
Ackr3
Rgmb
Itgb5
Fzd3
Chid1
Ligand EC
Receptor NRP
RJV
EC Ligand
NRP Receptor
logFC
4.8
-4.8
Il4
Tgfb3
Fgf1
Nampt
Cxcl12
Serpine1
Bmp4
Bdnf
Gdf11
Vegfc
Ccl2
Angpt2
Tnf
Ccl5
Ackr3
Acvr1b
Chid1
Hspg2
Rgmb
Vtn
Acvr2a
Ifnar1
Enpp2
Fgfr2
Cd44
Acvr1
Bambi
Itgb8
Lifr
Nr3c1
Itgav
Ifngr1
Adam9
Il10ra
Il15ra
Itgb1
Fzd1
Sorcs2
Mag
Ackr1
Nrp1
Grm3
Nr4a1
Bmpr1b
Ligand EC
Receptor NRP
AGA
EC Ligand
NRP Receptor
padj
> 0.05
0.01 to 0.05
0.001 to 0.01
0.0001 to 0.001
< 0.0001
logFC
3.6
-3.5
Serpine1
Bdnf
Ccl5
Angpt2
Il18
Nampt
Fgf1
Gdf11
Bmp4
Il4
Tgfb3
Nr4a1
Grm3
Sorcs2
Mag
Ackr1
Grm4
Ppara
Ahr
Grm5
Fzd9
Thbd
F2
Lrp1
Sdc4
Nr3c1
Itgav
Acvr2b
Acvr1b
Ltbp1
Grm2
Tgfbr1
Eng
Amhr2
Il15ra
Acvr2a
Cd44
Ifngr1
Rgmb
Il6st
Enpp2
Bambi
Itgb5
Fzd3
Chid1
Ligand EC
Receptor NRP

**Supplementary Fig 10. Intercellular communication networks between EC-OLG and EC-NRP revealed aging-related interactions that were modified by heterochronic parabiosis.** Canonical EC ligands and their cognate receptors in OLG (a) or in NRP (b) are shown in each paradigm (Aging, RJV, AGA). In all panels of ligand-receptor interactions, node color represents the magnitude of the DGE (logFC as estimated by DGE) such that the most significantly up-regulated genes are in magenta, and the downregulated genes are in blue. Node borders indicate FDR statistical significance of DGE. Edge color represents the sum of scaled differential expression magnitudes from each contributing node, while width and transparency are determined by the magnitude of the scaled differential expression (see details in the Methods section).

#### Supplementary Tables

**Supplementary Table 1. List of abbreviations for all cell types and subpopulations.**

**Supplementary Table 2. Metrics of pairwise comparison of cell types by cell number.** For each pairwise comparison, per cell type, ANOVA was applied, with significance designation (ns padj > 0.05, * padj ≤ 0.05, ** padj ≤ 0.01, *** padj ≤ 0.001).

**Supplementary Tables 3-11. DGE metrics per comparison, per cell type.** edgeR/muscat metrics were computed for each cluster in a comparison, with logFC, p-value, Benjamini-Hochberg p-value adjustment (p_adj.loc is per cluster, p_adj.global is with respect to all clusters in class) reported. TPM values ("tpm") for each animal type, per cell type are listed. Percentage expression ("pct") of each gene for each animal type, per cell type are listed. **RJV**=Rejuvenation-association ((OY-OX)-(OO-YX)), **AGA**=Aging Acceleration-associated ((YO-YX)-(YY-OX)), **Aging**=OX/YX, **OY_OO**=OY/OO, **OY_OX**=OY/OX, **OO_OX**=OO/OX, **YO_YY**=YO/YY, **YO_YX**=YO/YX, **YY_YX**=YY/YX. HypEPC and TNC clusters did not have enough cells distributed across multiple animals for inclusion in the DGE framework, but TPM and PCT are reported.

**Supplementary Table 12. RJV and Aging common DGEs and unique DGEs, per cell type**. DGE FDR ≤ 0.05 genes common between RJV and Aging, but also those only found in RJV, and those only found in Aging are listed per cell type.

**Supplementary Table 13. AGA and Aging common DGEs and unique DGEs, per cell type**. DGE FDR ≤ 0.05 genes common between AGA and Aging, but also those only found in AGA, and those only found in Aging are listed per cell type.

**Supplementary Table 14. DGE bi-directional DGEs between RJV and AGA.** For RJV and AGA DGE genes FDR ≤ 0.05 per cell type, report the genes that are RJV Up (logFC >0) and AGA Down (logFC<0), or RJV Down (logFC <0) and AGA Up (logFC>0). Count is the number of times a gene is bi-directionally expressed across cell types.

**Supplementary Table 15. Matrices of all DGE FDR ≤ 0.05 logFC values across cell types.** Per comparison, per gene, clusters where the gene's significance is FDR ≤ 0.05 have their logFC value reported. "Up" column is the sum of clusters with logFC >0, "Down" column is the sum of clusters with logFC < 0.

**Supplementary Table 16. DGE logFC values across all cell types**. Per comparison, per gene, collated logFC values across all clusters reporting DGE, with no thresholding. "Up" column is the sum of cell types with logFC >0, "Down" column is the sum of clusters with logFC < 0.

**Supplementary Table 17. Matrices of al identified significant GSEA terms per comparison across cell types**. fGSEA Benjamini-Hochberg adjusted p-value ≤ 0.25 significant terms are collated per comparison across all cell types by Normalized Enrichment Score (NES). Pathway and process metaclasses are described in the Methods. "Up" column is the sum of cell types with NES >0, "Down" column is the sum of cell types with NES < 0.

**Supplementary Table 18. SCENIC regulon matrices per animal type**. Per animal type, per cell type, SCENIC regulon activity scores are reported. Column "counts" is the sum of cell type that have a regulon score.

**Supplementary Table 19. Cell-cell communication networks in RJV, AGA, and per animal type.** For RJV, the set of unique source:target:receptor:ligand pairs that are found only in OY and YX combined. For AGA, the set of unique source:target:receptor:ligand pairs that are found only in YO and OX combined. YX, YY, YO, OX, OO, OY display the CellChat[7] networks derived for each animal type (see details in Methods). "Source" is the cell type the ligand comes from, while “Target” is the cell type found matching the ligand’s receptor. Probability and p-value are the statistical measures derived by CellChat. Ligand-Receptor pairs are given Interaction Names and assigned to a Pathway. Annotation provides the type of interaction are Secreted Signaling, ECM-Receptor, Cell-Cell Contact. Evidence codes and relevant PMIDs are provided by CellChat.

**Supplementary Table 20: Literature-curated senescence-associated genes**. Senescence-associated genes were curated from the literature for use as a reference gene set to perform fGSEA. HGNC.symbol denotes *Homo sapiens* gene symbol, MGI.ID denotes MGI ID number, MGI.symbol is *Mus musculus* gene symbol, Name is long name of the gene, and Feature Type denotes the type of gene or pseudogene.

**Supplementary Table 21: Senescence status GSEA per comparison.** For RJV, AGA, and aging, fGSEA against a literature-curated senescence gene set (Supplementary Table 20) to derive enrichment score, Normalized Enrichment Score (NES), p-value, Benjamini-Hochberg adjusted p-value, rank, and genes in the leading edge.
